# Callosal inputs generate side-invariant receptive fields in the barrel cortex

**DOI:** 10.1101/2022.11.17.516910

**Authors:** Roberto Montanari, Alicia Alonso-Andrés, Jorge Cabrera-Moreno, Javier Alegre-Cortés, Ramón Reig

## Abstract

Barrel cortex integrates contra- and ipsilateral whiskers’ inputs. While contralateral inputs depend on the thalamocortical innervation, ipsilateral ones are thought to rely on callosal axons. These are more abundant in the barrel cortex region bordering with S2 and containing the row A-whiskers representation, the row lying nearest to the facial midline. Here we ask what role this callosal axonal arrangement plays in ipsilateral tactile signalling. We found that novel object exploration with ipsilateral whiskers confines c-Fos expression within the highly callosal subregion. Targeting this area with *in vivo* patch-clamp recordings revealed neurons with uniquely strong ipsilateral responses dependent on the corpus callosum, as assessed by tetrodotoxin silencing and by optogenetic activation of the contralateral hemisphere. Still, in this area, stimulation of contra- or ipsilateral row A-whiskers evoked an indistinguishable response in some neurons, mostly located in layers 5/6, indicating their involvement in the midline representation of the whiskers’ sensory space.

## Introduction

The corpus callosum (CC) is the characteristic commissure of placental mammals’ brains^1^. The CC is mainly composed of glutamatergic axons interconnecting the two hemispheres and belonging to neocortical pyramidal neurons^2,3^. Besides allowing the transfer of simple discriminative memories from the trained to the untrained hemisphere^4^, the CC participates in motor and sensory operations. More specifically, in primary visual and somatosensory cortex (S1), it is involved in the fusion at the midline of the right and left sensory hemi-spaces^5–9^. Accordingly, neurons dedicated to the midline of the visual and somatosensory space exhibit a side-invariant receptive field (RF). Namely, they manifest comparable sensitivity to the stimulation of sensory organs on either side of the body. This effect is obtained through the CC that replicates the primary thalamocortical response of the activated hemisphere into the opposite hemisphere, thus generating in the latter a mirror replica of the activation that occurred in the former^6^. By generating side-invariant RFs, midline-crossing circuits would guarantee a continuity between the right and left sensory maps^9,10^. Congruently with this binding role, in S1 of placental mammals, side-invariant neurons represent midline and para-midline body regions like nose, chin, intraoral surfaces, ventral/dorsal trunk etc., and often occupy cytoarchitectonic borders of neocortical areas^6^. On the contrary, in representations of body parts far from the midline (e.g., extremities) callosal connections are scant or absent (but see ref.^7^).

Rodent whiskers are somatotopically represented in the barrel cortex (BC). Thalamic innervation enables BC neurons to respond to contralateral whisker stimulations, while ipsilateral responses are commonly attributed to callosal innervation given the complete decussation of the trigeminothalamic tract^11–16^. In rodents’ BC, callosal innervation is scant and involves supra- and infragranular layers, for both axonal origin and termination^17^. Axons entering from the white matter ascend towards the neocortical surface through septal dysgranular areas, extensively avoiding layer (L) 4 barrel hollows^18,19^. The densest innervation is restricted to a narrow stripe at the S1/S2 cytoarchitectonic border^18–25^. In the BC, this border contains the representation of row A-whiskers, the row that lies nearest to the facial midline^22,23^. This resembles the S1 representation of other midline and para-midline body parts and led us to ask 1) how the interhemispheric synaptic transmission varies across distant regions of the BC that display different degrees of CC innervation and 2) whether or not the whiskers system also shows side-invariant RFs obeying the midline rule.

Our strategy involved testing the functional relevance of callosal axons’ disposition in the BC by c-Fos activation in mice exploring novel objects with ipsilateral whiskers inputs only. Then, through *in vivo* whole-cell patch-clamp we compared whisker responses in distal regions of the BC representing either para-midline (row A) or lateral (row E) whiskers. We found stronger and faster ipsilateral responses upon row A stimulation that were produced by the presynaptic excitation generated by the opposite hemisphere through the CC. Lastly, we report the presence of side-invariant responses dependent on the CC, which demonstrates that the whiskers system conforms to the midline rule.

## Results

### Ipsilateral whiskers induce regionalised c-Fos expression in the barrel cortex upon exploration

Whiskers-mediated tactile experience drives the somatotopic expression of c-Fos in the BC^26^, indicating the metabolic activation of whiskers-dedicated neurons. The BC can respond to the stimulation of ipsilateral whiskers^11–16^, but how the arrangement of callosal innervation affects its recruitment is unknown. First, we studied the arrangement of callosal projections in the BC by biotin dextran amide (BDA) injections in the opposite BC’. Injections restricted to the posterolateral aspect (i.e., row A/B territory) of the BC showed higher axonal density at homotopic locations in the opposite BC’ than injections restricted to the anteromedial aspect (i.e., row D/E territory) (Fig. 1a–c). This result confirmed the patterned disposition of callosal axons connecting the BCs. To explore the functional significance of this disposition, we extracted the density of c-Fos^+^ expression in the BC of mice that were subjected to manipulations of their whisker arrays and to a subsequent period of free arena exploration with novel objects. Density of c-Fos^+^ nuclei was quantified on flattened cortical preparations showing barrels’ outlines (see Methods). To avoid confusing the thalamocortical activity evoked by the contralateral whiskers with that evoked by the ipsilateral whiskers, we first looked for a method to isolate the BC from peripheral inputs of both sides. Hence, we trimmed the whiskers bilaterally at their base (Fig. 1e; n_hem_=10, 2 hemispheres/mouse). However, the density of c-Fos^+^ nuclei across the BCs in this condition (Sup. Tab. 1) did not differ from the control condition in which mice (Fig. 1d; n_hem_=6, 2 hemispheres/mouse) retained the entire set of bilateral whiskers (Fig. 1h; p=0.3132). This suggested that, even with characteristic patterns (Sup. Fig. 1a,b), BCs were still recruited despite the complete trimming of whiskers. Since trimming whiskers could not abolish the BC activation, the method could not be used to isolate the ipsilateral component of the BC activity. Subcutaneous injection of lidocaine in the whisker-pad is a previously described method for long-lasting suppression of whisker responses in the BC^27^. Indeed, in mice receiving lidocaine bilaterally (n_hem_ =8, 2 hemispheres/mouse), BCs were nearly devoid of c-Fos expression (Fig. 1f), displaying a very significant reduction compared to the trimmed whiskers condition (Fig. 1h; p=4.5·10^-5^). This indicated, first, that BCs could not receive enough peripheral input to trigger c-Fos expression and, second, that no other brain areas (e.g., primary motor cortex) could do it either. Finally, we injected lidocaine unilaterally and measured c-Fos positive cell density in the BC contralateral to the injected side, which was deprived of direct thalamocortical input (input-blocked BC’; Fig. 1g; n_hem_=4; 1 hemisphere/mouse). Density was higher in the input-blocked BC’ than in BCs sensory-deprived through bilateral lidocaine injections (Fig. 1h; p=0.0162). This suggested that the opposite, input-receiving BC triggered activity in the input-blocked BC’ through the callosal innervation. To confirm a callosal activation, we checked for anatomical overlap between c-Fos density and callosal fibres, hence we normalised the density and assigned it to the different whiskers rows. Whereas c-Fos density was homogeneously distributed across rows in the input-receiving BC (p= 0.5625), we found that was unevenly distributed in the input-blocked BC’ (p= 0.0021), where it concentrated in the CC-recipient posterolateral aspect (i.e., row A- and, to a lesser extent, row B-barrels; Fig. 1i). In input-blocked BC’, CC-recipient septa also showed a strong, yet non-significant trend of higher posterolateral c-Fos density (Fig. 1j; p=0.0507). These results demonstrate that, during novel object exploration, the posterolateral aspect of the BC’ is the main recipient of the activity raised by ipsilateral whiskers, which is likely recruited by the callosal pathway.

**Fig. 1.**
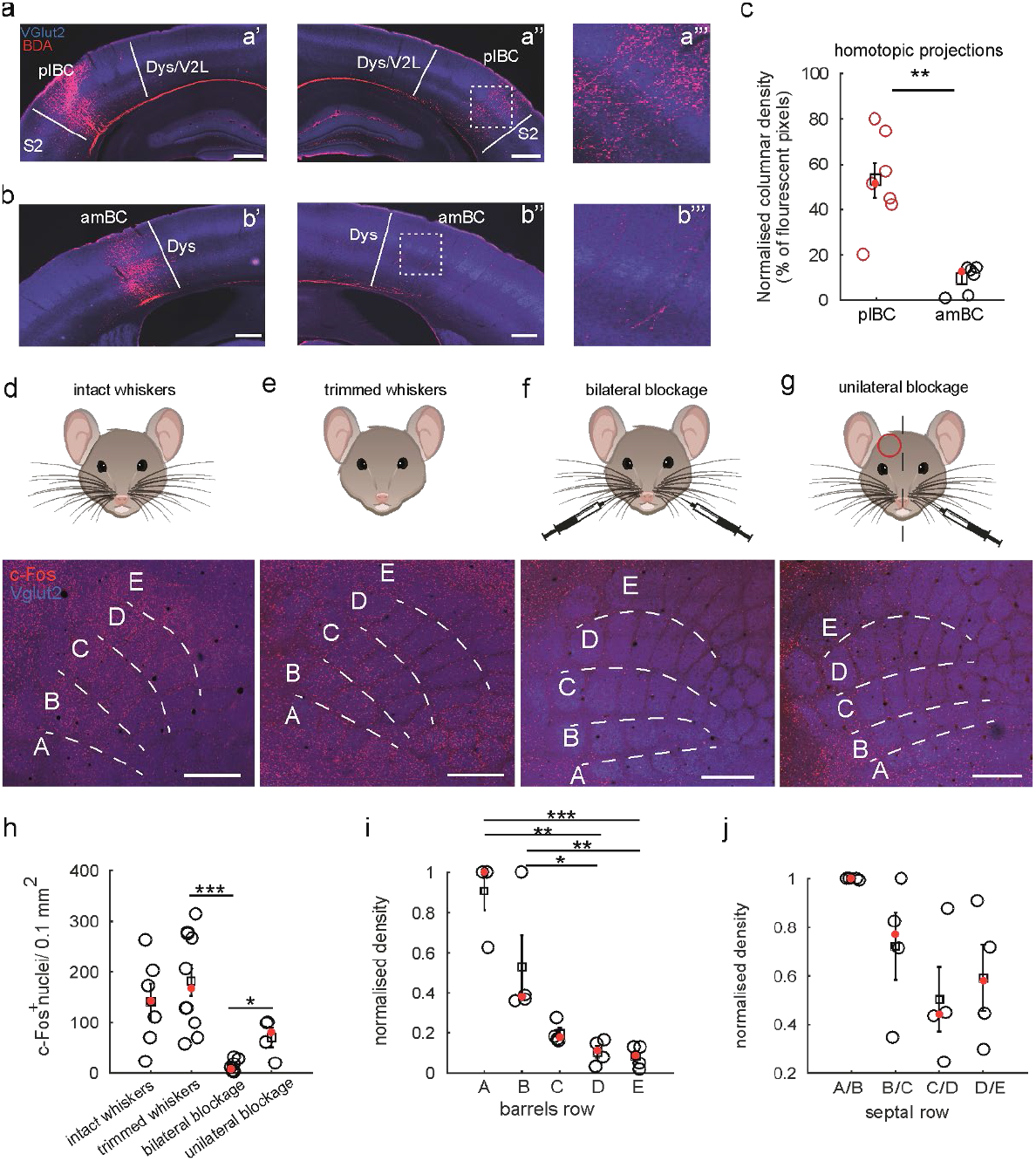
Callosal innervation and the recruitment of c-Fos expression in the barrel cortex. **(a)** BDA injection in plBC (a’) with contralateral callosal innervation of the homotopic plBC (a’’). a’’’ shows inset from dashed square in a’’. **(b)** BDA injection in amBC (b’) with contralateral callosal innervation of the homotopic amBC (b’’). b’’’ shows inset from dashed square in b’’. **(c)** Quantification of homotopic projections. **(d-g)** Cartoon of the experimental paradigm (up) and c-Fos expression (bottom). **(d)** Mice carrying bilaterally intact arrays of whiskers. **(e)** Bilaterally trimmed whiskers mice. Note, the BC expresses c-Fos despite the peripheral intervention. **(f)** Bilateral input block through injections of lidocaine in the whisker-pads nearly abolishes c-Fos expression in the BCs. **(g)** Expression regionalises towards row A and B-whiskers in the BC upon unilateral input blockage of the contralateral whisker-pad. Red circle indicates the side of input-blocked BC’. Images obtained by Z-projection across the tissue depth of maximal intensity for both channels. **(h)** Statistical comparisons of the density of c-Fos^+^ nuclei in the different conditions. (**i**) Comparison of c-Fos expression across barrel rows normalised by the maximum density in each experiment (n=4; KW density input-receiving BC: df=4, *χ*^2^=2.97, p=0.5625; KW density input-blocked BC’: df=4, *χ*^2^=16.86, p=0.0021; Fisher’s LSD post-hoc density input-blocked BC’: p_A vs B_= 0.5093; p_A vs C_= 0.0673; p_A vs D_= 0.0030; p_A vs E_= 7·10^-4^; p_B vs C_= 0.2421; p_B vs D_= 0.0209; p_B vs E_= 0.0063; p_C vs D_= 0.2544; p_C vs E_= 0.1188; p_D vs E_= 0.6746). (**j**) Same as i for septa between barrel rows (n=4; KW density input-blocked BC’: df=3, *χ*^2^ =7.79, p=0.0507). Dys= dysgranular area; Dys/V2L= dysgranular area/secondary visual cortex; S2 = secondary somatosensory cortex. Scale bars: a-b= 500 μm, c-f= 400 μm.

### Laminar postsynaptic responses to contra- and ipsilateral whisker stimulation in the barrel cortex

Since the posterolateral BC subregion receiving high callosal innervation remained active despite the unilateral block of whiskers input, while less innervated areas were accordingly less active (Fig. 1f,h), we characterised the corresponding contra- and ipsilateral whisker responses. In order to observe whisker responses in the areas of interest, we performed *in vivo* whole-cell patch-clamp recordings in anaesthetised mice. Craniotomies were made following coordinates in ref.^28^, targeting the representation of row A and row E barrel hollows and sides (Fig. 2a,c), representing the whiskers nearest and farthest from the facial midline, respectively. Prior to whole-cell recordings, we performed a whiskers-to-barrels mapping by recording LFP responses upon contralateral whiskers stimulation matching the target cortical representation (Sup. Fig. 3c), here referred to as somatotopic stimulation. To further assess the targeting of selected areas, electrode tracks and recorded neurons were anatomically localised by biocytin-streptavidin staining in the corresponding brain sections (Fig. 2c). Considering that few unrecovered neurons were also included in the analysis, we refer to the row A-containing area to as posterolateral BC (plBC), and to the row E-containing area to as the anteromedial BC (amBC). Groups of whiskers (either A_2_-A_3_ or E_2_–E_3_) were stimulated with a custom-made, solenoid-based whisker puller inspired by ref.^29^ (Sup. Fig. 3a,b). Stimulations consisted of a firm caudo-rostral displacement [force: 7.4 ± 1.8 mN (mean ± s.d.); travel: 3.5 mm; duration: 15 ms] of the whiskers glued to the tip of a stainless-steel ultrafine wire, sliding backwards upon solenoid electrification. Once a cell was patched, somatotopic contra- and ipsilateral stimulations were delivered every 3 or 5 seconds (Fig. 2a,b). Under anaesthesia, the membrane potential (V_m_) of cortical neurons oscillates periodically at ~1 Hz, alternating between up- and down-states (Fig. 2b). Because these spontaneous network transitions have a strong impact on whisker responses^30,31^, we analysed separately stimulations/responses falling in either up- or down-state in 18 L2/3 neurons, 15 L4 neurons and 37 L5/6 neurons.

**Fig. 2.**
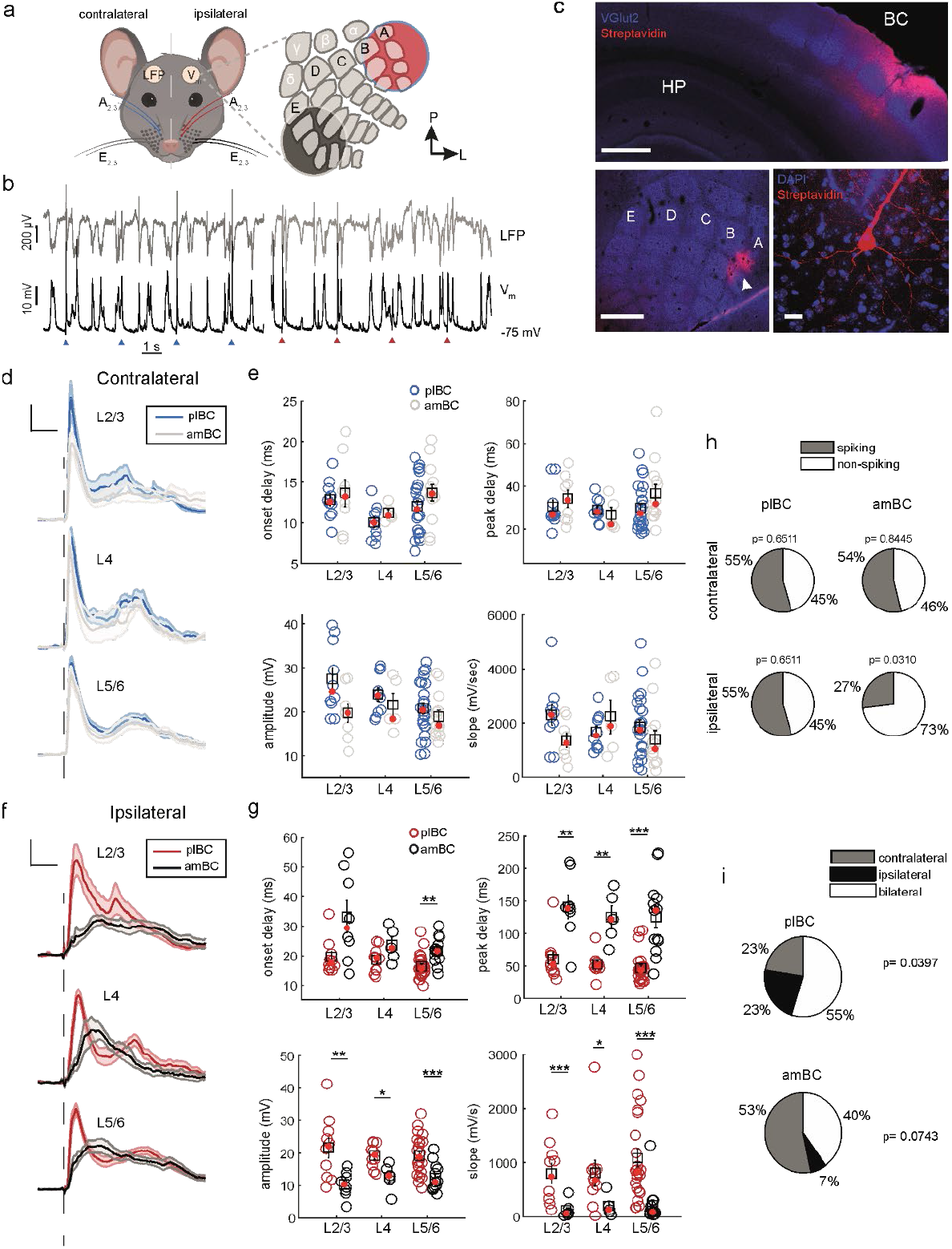
Ipsilateral whisker responses are more vigorous in the posterolateral barrel cortex. **(a)** Cartoon of sides and sets of stimulated whiskers, colour-coded according to the cortical subregion recorded, shown on the right. **(b)** Example of recordings in the posterolateral barrel cortex with whiskers stimulation. Blue and red arrowheads for contra- and ipsilateral stimulations, respectively. **(c)** Immunohistochemistry for assessing the placement of the whole-cell patch-clamp recording and cell anatomy. Upper image: example coronal section of a posterolateral barrel cortex recording site with biocytin-streptavidin staining. Scale bar= 400 μm, BC= barrel cortex, HP= hippocampus; Lower images: on the left, example flatten brain preparation of L4 with biocytin-streptavidin staining within the row A representation (white arrowhead). Scale bar= 400 μm; on the right, example of a recovered pyramidal neuron with biocytin-streptavidin staining (red), Vglut2 staining (blue). Scale bar= 20 μm; **(d)** Grand average response ± S.E.M (shadows) following somatotopic contralateral stimulations along the cortical depth, aligned at stimulus delivery time (dashed line). **(e)** Statistical comparison of parameters of responses in d. **(f)** Same as d for the same pool of neurons responding to somatotopic ipsilateral stimulations. **(g)** Statistical comparison of parameters of responses in f. **(h)** Proportions of spiking and non-spiking neurons in the four conditions of stimulation/recording (for df=1 in contralateral, plBC: 24/44, *χ*^2^=0.2045; amBC: 14/26, *χ*^2^=0.0385; for df=1 in ipsilateral, plBC: 24/44, *χ*^2^=0.2045; amBC= 7/26, *χ*^2^=4.6538). **(i)** Distribution of suprathreshold receptive fields in spike-responding neurons (for df=2, plBC: *χ*^2^=6.4516; amBC: *χ*^2^=5.2000).

Firstly, we compared between ipsi- and contralateral evoked responses during down-state for all recorded neurons. Ipsilateral stimulations evoked responses characterized by longer onset delays [(ms): contra.=12.3 (9.1-14.3), ipsi.=18.6 (15.4-23.5), p=2·10^-5^], weaker amplitude [(mV): contra.=20.2 (16.9-26.8), ipsi.=15.9 (11.6-21.9), p=5·10^-15^] and lower probability to evoke action potentials compared to their contralateral counterparts [P(AP): contra.= 0.67 (0.2-1), ipsi.= 0.16 (0.04-0.5), p=5·10^-4^].

Then, we studied how neurons of different layers responded to contra- and ipsilateral whisker deflections. Contralateral displacement of the whiskers robustly evoked responses, with the thalamorecipient L4 [10.7 (9.2-11.4) ms] showing the fastest onset delay, followed by L2/3 [12.6 (11.1-14.5) ms] and L5/6 [13.1 (8.8-14.7) ms] (KW: df=2, *χ*2=6.44, p=0.040). In the same neurons, ipsilateral stimulation evoked responses in which onset delay of L5/6 tended to precede upper ones in a nearly significant fashion [L5/6: 16.5 (14.5-21.4) ms, L4: 19.6 (15.6-24.1) ms, L2/3: 19.6 (16.6-32.7); KW: df=2, *χ*^2^=4.792, p=0.091]. Indeed, if recording depth was considered instead of laminar identity, there was a significant inverse correlation with onset delay, with deeper neurons responding faster than upper ones (Pearson’s pairwise corr.: r=0.36, p=0.0018). Consistent with other descriptions^6,12,14–16^, these data indicate that longer delay, lower amplitude, reduced spiking, and a different pattern of columnar activation distinguish S1 ipsilateral responses from contralateral ones.

### Ipsilateral responses are stronger in the posterolateral barrel cortex

Next, in order to relate ipsilateral responses to distinct callosal innervation patterns, we compared neurons recorded in plBC (n=44) and amBC (n=26) in the different layers. Contralateral responses, except L2/3 neurons that tended to show higher amplitude in plBC than amBC (only significant during up-state; Sup. Fig. 2b,d), did not differ in onset and peak delay, amplitude, and slope across layers (Fig. 2d,e and Sup. Tab. 2). By contrast, ipsilateral responses differed in most of these aspects between the same plBC and amBC neurons. In all layers, responses had a significantly faster peak, higher amplitude, and steeper slope in plBC than in amBC neurons, with L5/6 response onset significantly faster in plBC than in amBC ones (Fig. 2f,g and Sup. Tab. 2). In addition, during up-state, ipsilateral stimulation elicited responses in the majority of plBC neurons, but rarely in amBC neurons (Sup. Fig. 2a,c). Contrary to subthreshold responses that could be evoked in all the neurons independently of the stimulation side during down-sate, only a fraction showed a suprathreshold response. Indeed, following contralateral stimulation, spike-responding neurons in both plBC and amBC represented roughly half of the pool (Fig. 2h). Interestingly, following ipsilateral stimulation, the presence of spike-responding neurons decreased in amBC but not in plBC (Fig. 2h), suggesting an increased ipsilateral output in the latter. We also analysed the laminar distribution of spike-responding neurons upon ipsilateral stimulation (Fig. 2h). While spike-responding neurons were evenly dispersed throughout layers in amBC (df=2, *χ*^2^= 2, p= 0.3679), they were significantly concentrated in L5/6 in plBC (15/19, df=2, *χ*^2^= 9.25, p= 0.0098), suggesting downstream targets being preferentially subcortical or interhemispheric. Considered across layers, suprathreshold RFs (Fig. 2i) in plBC were most commonly bilateral (17/31, 54%), with remaining neurons equally split between contra-(7/31, 23%) and ipsilateral RFs (7/31, 23%). Ipsilateral suprathreshold RFs were almost absent in amBC (1/15, 7%), with a nearly significant tendency for contralateral RFs to be the most represented class (8/15, 53%), followed by the bilateral one (6/15, 40%). Hence, while the plBC seems to equally represent both the ipsi- and contralateral sensory hemi-spaces, the amBC is mostly dedicated to the contralateral one. The greater ipsilateral sensitivity of plBC neurons compared to amBC neurons could not be attributed to different intrinsic excitability, since their electrophysiological membrane properties were not different in either up- or down-states (Sup. Tab. 3), instead suggesting a distinct synaptic recruitment. Overall, these results demonstrate that the plBC is populated by neurons endowed with a RF highly sensitive to the stimulation of ipsilateral (row A) whiskers, likely due to stronger callosal inputs.

### Ipsilateral responses are controlled by feed-forward inhibition

In the BC, feed-forward inhibition is best known to act at thalamocortical synapses through a disynaptic mechanism mediated by Parvalbumin-positive or Somatostatin-positive GABAergic interneurons^33,34^. This type of inhibition can modulate the time window for thalamic input integration^35^ and be governed by the direction of whisker deflections^36^. Like thalamocortical afferents, callosal afferents can also mediate disynaptic feed-forward inhibition in distinct neocortical areas of rodents^37–39^ and cats^3^. Yet, despite the presence of direct callosal contacts on inhibitory interneurons^40^ and the increased firing of putative fast-spiking neurons upon ipsilateral stimulation^16^, to our knowledge no study so far has directly demonstrated the presence of inhibitory ipsilateral responses elicited by feed-forward inhibition in the mouse S1. To address that, and in order to decompose the excitatory and inhibitory response components, we used a low-chloride intracellular solution (see Methods), enabling us to hold recorded neurons at the reversal potential for excitation (~ −5 mV) or GABAa inhibition (~ -70 mV). By injecting positive currents to depolarise neurons close to their excitatory reversal potential, we sampled inhibitory responses to contra-(n=7) and ipsilateral (n=19) somatotopic whisker stimulation.

As expected, contralateral stimulations could reliably evoke inhibitory responses (Fig. 3a), whose peak delay was nearly coincident with the excitatory one of the entire neuronal pool [contra. peak delay (ms): exc=27.8 (24.2-37.8), n=70, inh=30.2 (29.2-36.7), n= 7, p=0.2568]. Comparable to their contralateral counterparts, ipsilateral stimulations also elicited responses with the excitatory and inhibitory components (Fig. 3a,b). The amplitude of inhibitory and excitatory ipsilateral responses was significantly correlated when both were recorded in the same neurons (n= 14; Fig 3b), with inhibitory response amplitude being comparatively smaller. To observe whether ipsilateral inhibitory responses could vary similarly to the excitatory ones following the extent of local callosal innervation (Fig. 1a,b), we compared plBC neurons (n=10) with amBC ones (n=9) (Fig. 3c). As with the excitatory component, inhibition was faster and much more vigorous in plBC than in amBC. Indeed, they had shorter onsets and peak delays, higher amplitudes, and steeper slopes in plBC than in amBC neurons (Fig 3d & Sup. Tab. 4), suggesting that the inhibitory responses also relied on callosal innervation. Furthermore, both the excitatory and inhibitory components seemed to show different temporal dynamics between the two subregions. Therefore, we compared the onset and peak delay separately for plBC and amBC. In both subregions, the onset of inhibitory responses was significantly delayed with respect to excitatory responses, consistent with a feed-forward inhibition mechanism. Yet, this was accompanied by regional differences underlying the onset of the inhibitory component. Specifically, in plBC the inhibitory component was delayed only by ~5 ms compared to the excitatory one (medians’ difference), while in amBC by ~12 ms [Fig. 3d,e; onset (ms): plBC: E=16.4 (14.8-19.5), I=21.3 (19.3-27.8), p=0.0010; amBC: E=22.5 (19.1-28.1), I=34.3 (32.9-44.1), p=0.0008], suggesting the involvement of different pathways in the generation of inhibitory responses. These results indicate that ipsilateral whisker responses are controlled by feed-forward inhibition, whose strength and temporal dynamics vary with the amount of excitation received and the extent of local callosal innervation.

**Fig. 3.**
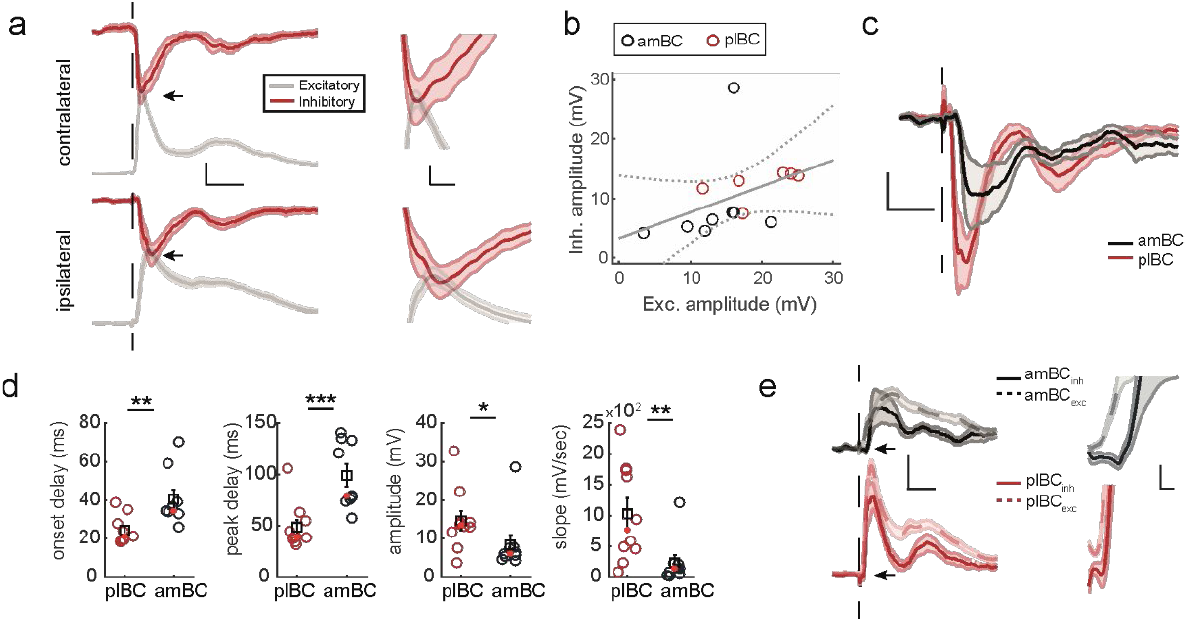
Ipsilateral whiskers stimulation recruits feed-forward inhibition in the barrel cortex. **(a)** Left: grand average ± S.E.M (shadow) of excitatory (gray) and inhibitory (red) responses aligned at stimulus delivery time (dashed line). Vertical bar=5 mV, horizontal bar=100 ms. Right: insets from black arrows on the left showing the nearly coincident peak between excitation and inhibition in both contra- and ipsilateral responses. Vertical bar=2 mV, horizontal bar=10 ms. **(b)** Inhibitory amplitude as a function of excitatory amplitude in response to somatotopic whiskers stimulation in the same neurons [exc.-inh. correlation (Spearman), all neurons considered together: rho= 0.62, p= 0.0202]. **(c)** Grand average ± S.E.M (shadow) of inhibitory responses aligned at stimulus delivery time (dashed line) recorded in the two barrel cortex subregions. Vertical bar=5 mV, horizontal bar=100 ms. **(d)** Statistical comparison of parameters for responses in c. **(e)** Left: excitatory (dashed line) and inhibitory (solid line) grand averages ± S.E.M (shadow) in the two BC subregions aligned at stimulus delivery time (vertical dashed line), with inhibitory responses inverted in sign with respect to c for visualisation. Vertical bar=5 mV, horizontal bar=100 ms. Right: insets from black arrows on the left showing the grand average around the onset of the excitatory and inhibitory components. Vertical bar=1 mV, horizontal bar=10 ms.

### Ipsilateral responses in the posterolateral barrel cortex have their origin in the contralateral hemisphere

What is the source of the vigorous ipsilateral response? While normally described as totally decussating towards the contralateral somatosensory thalamus^14^, principal sensory nucleus (PrV) axons branch extensively^41^, and some of them, emanating from its dorsal portion, reach the ipsilateral thalamus^42^. The dorsal PrV represents midline body parts such as the lower jaw and lips^43^, with the representation of the former also receiving CC afferents at the cortical level^44^. Although row A-whiskers are represented in the ventral PrV, they lie next to the facial midline and cortically receive CC afferents. Thus, we could not exclude *a priori* that few ipsilateral trigeminothalamic axons, or an earlier commissure, could have passed undetected so far and be responsible for the vigorous ipsilateral response in plBC. To exclude these possibilities and learn about the contribution of callosal neurons to the sensory response, we injected tetrodotoxin (TTX 1-1.5 μL, 100 μM) in one BC while recording in the opposite plBC (n=11, 1 neuron/mouse) (Fig 4a). Contra- and ipsilateral row A-whiskers were stimulated before (CTRL) and after (TTX) the drug application, keeping the neuron under whole-cell patch-clamp. Amplitude of LFP downward deflections in the injected BC greatly decreased upon TTX application (Fig 4a,b), indicating a strong depression of its excitatory activity. In the opposite hemisphere, although a weak depolarisation could still be evoked, the response to ipsilateral row A-whiskers stimulation lost its timing and vigour (Fig 4c). Indeed, onset and peak delays elongated, the amplitude was strongly decreased, and responses rose significantly more slowly upon TTX application (Fig. 4d). In conclusion, ipsilateral responses lost vigour when contralateral activity was blocked. This suggests a weaker synaptic recruitment in TTX compared to CTRL, demonstrating a strong dependence of the ipsilateral response on callosal neurons of the opposite BC.

**Fig. 4.**
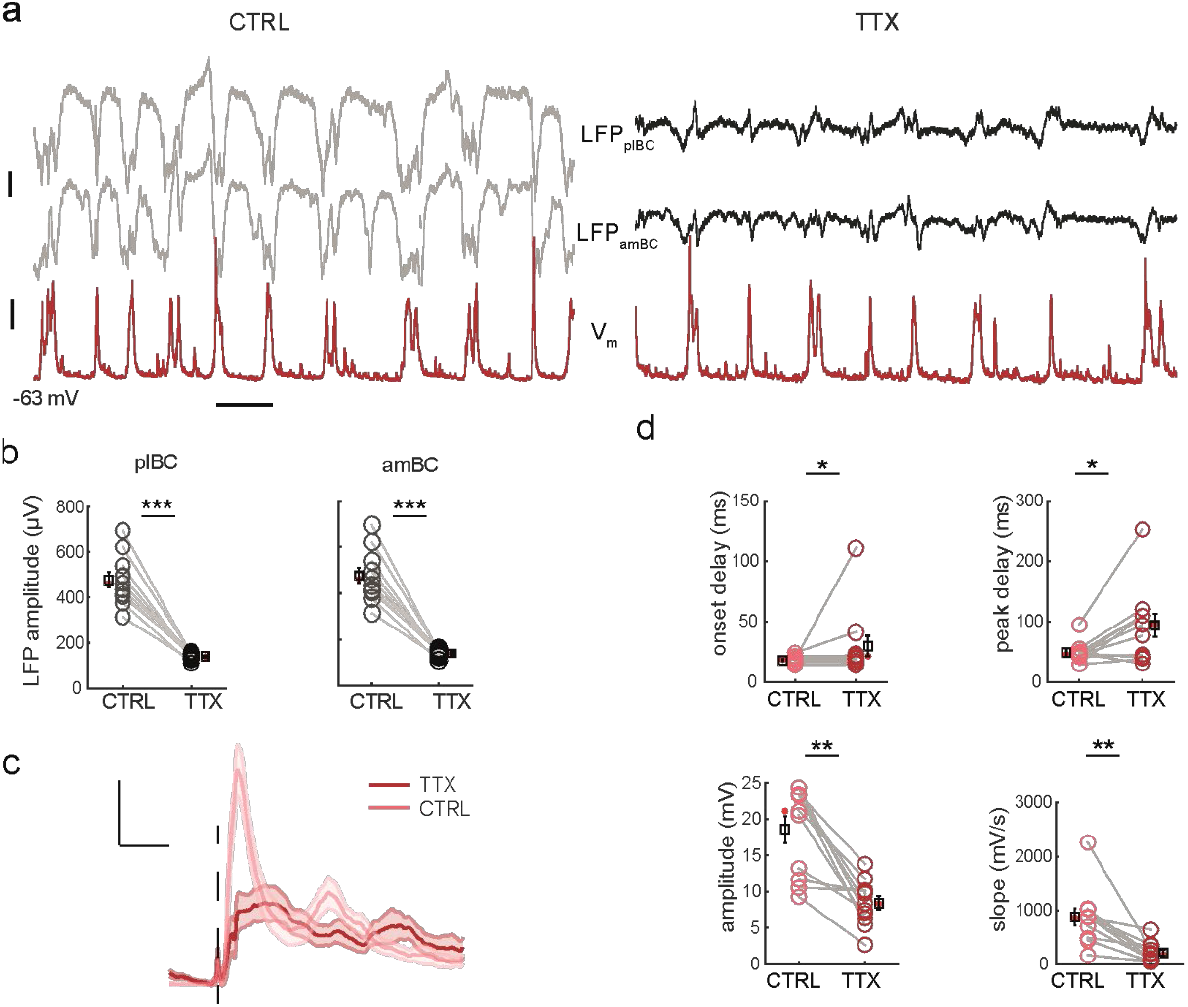
Synaptic recruitment deteriorates upon activity blockage of the opposite barrel cortex. **(a)** Example recording of a plBC neuron V_m_ (red bottom trace) and contralateral plBC and amBC LFPs (left, grey traces) during spontaneous slow-wave oscillations. Note the nearly complete activity suppression of the contralateral barrel cortex, at both plBC and amBC locations, upon TTX application (right, black traces). **(b)** Statistical comparison of LFP downward deflections’ amplitude between CTRL and TTX [plBC amplitude (μV): LFP_CTRL_= 463.0 (408.4-528.1), LFP_TTX_= 141.4 (130.7-152.6), WSR: p=9.7·10^-4^; amBC amplitude (μV): LFP_CTRL_= 418.0 (335.0-515.8), LFP_TTX_= 148.9 (137.9-171.3), WSR: p=8.5·10^-5^]. **(c)** Grand average ± S.E.M. (shadow) of row A-whiskers responses aligned at stimulus delivery time (dashed line) in the two conditions. Vertical bar= 5 mV, horizontal bar= 100 ms. **(d)** Statistical comparison of parameters of responses in c [WSR: onset delay (ms): CTRL=18.6 (15.7-19.4), TTX=21.3 (18.7-22.8), p=0.0313; peak delay (ms): CTRL=46.2 (42.8-49.1), TTX=94.5 (42.2-109.9), p=0.0137; amplitude (mV): CTRL=21.1 (12-23.3), TTX=8.3 (6.4-10.1), p=0.0010; slope (mV/s): CTRL=865 (538-985), TTX=178 (67.1-320), p=0.0010].

Yet, the persistence of a weak evoked response after contralateral TTX blockage required a further validation of the callosal scenario. If the CC was driving these responses, then directly stimulating the opposite BC should have evoked responses reflecting the difference in callosal innervation between plBC and amBC (Fig. 1a–c), similar to the sensory ones evoked from the whiskers (Fig. 2f). In this case, the residual response observed in TTX would be accounted for by an incomplete activity suppression of the drug-receiving hemisphere. To explore this issue, we took advantage of NEX-Cre mice that allow the genetic targeting of cortical projection neurons^45^. We injected a Cre-dependent AAV-ChR2 carrying EYFP in the BC to impose its excitation by blue-light illumination (5 or 10 ms light-step), while recording neurons in the opposite plBC (n=11) or amBC (n=12). To be sure about restricting the activation of only homotopic contralateral areas, for most of the recorded neurons (20/23) two LFPs were collected in amBC and plBC of the illuminated hemisphere, with the optic fibre positioned over one of the two, in mirror-symmetric (i.e., homotopic) opposition to the contralateral patch-clamp electrode (Fig. 5a,b).

**Fig. 5.**
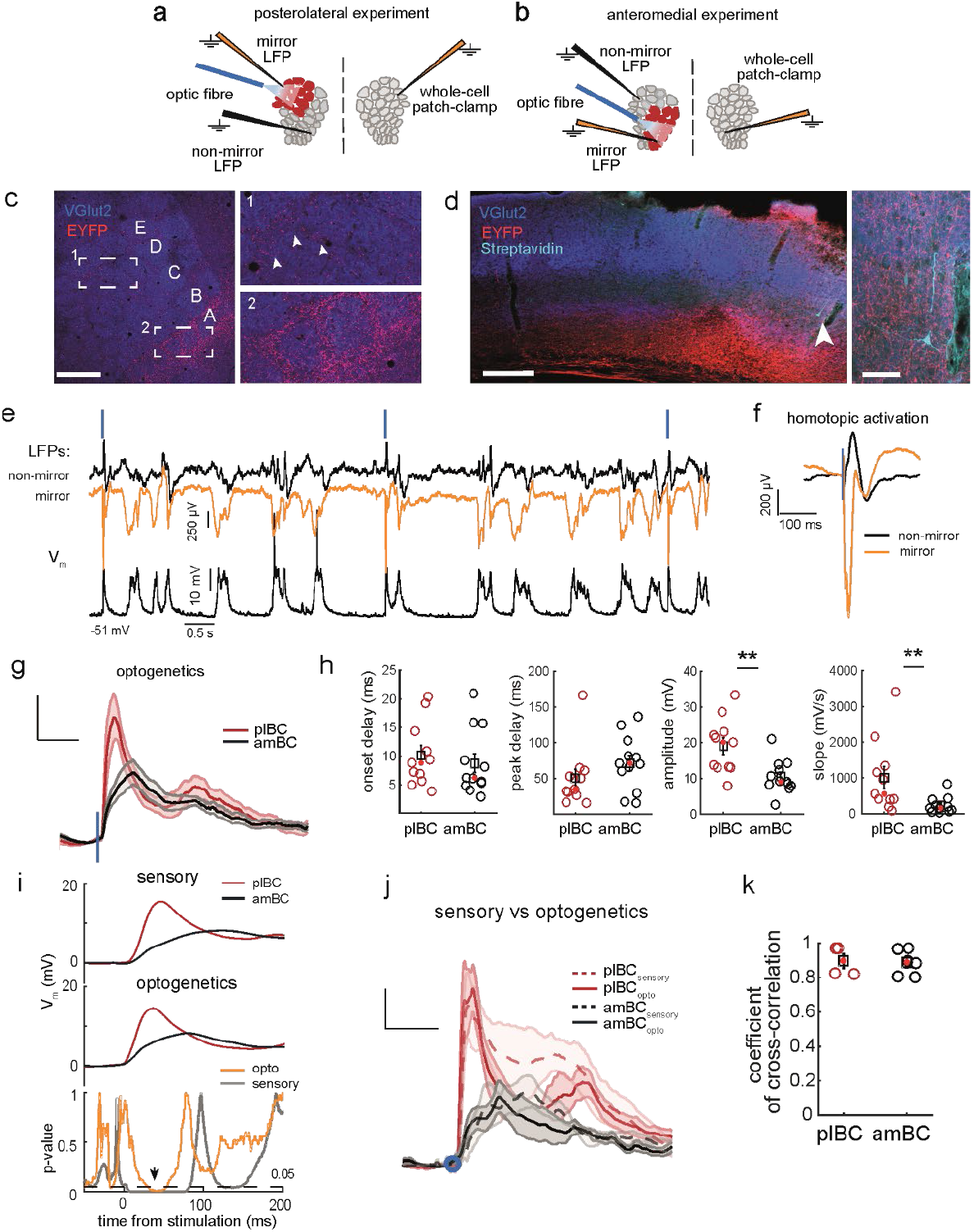
Homotopic cortically-evoked responses recapitulate sensory ones. **(a)** Schema of the experimental setting for homotopic optogenetics stimulation/recording in the plBC. **(b)** Same as a for the amBC. **(c)** Flatten L4 preparation of the barrel cortex contralateral to the AVV-EYFP-ChR2 injected hemisphere of an example NEX-Cre mouse. On the right, magnification from dashed white areas shown on the left. White arrowheads show a scanter callosal innervation of septa (top) compared to the row A representation (bottom). Scale bar= 500 μm. **(d)** Same animal preparation as c but in coronal section. Scale bar= 200 μm. White arrowhead indicates the recorded pyramidal neuron showed on the right within the bulk of invading callosal axons. Scale bar= 50 μm. **(e)** Example stimulation/recording of a homotopic optogenetics experiment in the plBC. Top traces LFPs, bottom trace V_m_. **(f)** Example waveform averages of mirror and non-mirror LFPs in homotopic experiments aligned at stimulus delivery time (blue line). **(g)** Grand average ± S.E.M. (shadow) of the whole-cell recordings with optogenetics responses aligned at stimulus delivery time (blue line) in the two subregions. **(h)** Statistical comparison of parameters of responses in g [onset delay (ms): plBC=8.8 (5.9-13.5), amBC=6.1 (5.0-12.5), p=0.4060; peak delay (ms): plBC=34.9 (28.1-59.0), amBC=71.1 (45.0-92.4), p=0.1029; amplitude (mV): plBC=20.1 (13.0-22.8), amBC=9.0 (8.1-13.0), p=0.0089; slope (mV/s): plBC=561.4 (397.4-1.3·10^3^), amBC=152.1 (80.5-309.0), p=0.006]. **(i)** Grand averages of sensory (top) and optogenetics (middle) responses of plBC and amBC neurons and p-value distributions (bottom) resulting from their comparison. Black arrow indicates the region of overlap for p < 0.05. **(j)** Grand average ± S.E.M. (shadow) of sensory and optogenetics responses recorded in the same neurons aligned at response onset (blue circle). **(k)** Cross-correlation between sensory and optogenetics responses of the neural populations inj [coeff._sensory/opto_.: plBC (n=4): 0.89 (0.82-0.97), amBC (n=6): 0.89 (0.80-0.96)].

Again, the great majority of EYFP-positive callosal axons invaded the BC at the S1/S2-border (Fig. 5c, d), enveloping the row A-whiskers representation with more scant labelling of the septa (Fig. 5c). Upon blue-light illumination, the mirror symmetric LFP activated locally and powerfully (Fig. 5e, f). Clearly reflecting a more vigorous input from the opposite hemisphere, optogenetic responses of plBC neurons had much higher amplitudes and steeper slopes than those of amBC, while the remaining parameters were similar (Fig. 5g, h). As we predicted, the overall difference in shape between amBC and plBC responses recapitulated with fidelity the difference in dynamics between the ipsilateral whiskers’ responses. Indeed, statistically comparing the waveforms between plBC and amBC neurons evoked by optogenetics and sensory stimulations resulted in a similar pattern of significant (and non-significant) p-values over time (Fig. 5i). Additionally, in neurons receiving both the ipsilateral whiskers and interhemispheric optogenetics stimulations (Fig. 5j; plBC n=4, amBC n=6), cross-correlation between waveforms of the two evoked responses was extremely high in both subregions (~0.90, Fig. 5k), likely reflecting the shared recruitment of CC fibres across stimulation types. These results strongly suggest that the callosal input alone underlies the different ipsilateral responses between plBC and amBC neurons.

### Ipsilateral responses mimic their contralateral counterparts in the posterolateral barrel cortex

Given the proximity to the facial midline of row A-whiskers, the callosal afferents invading its cortical representation at the S1/S2 border, and the vigorous callosal response reminiscent of a thalamocortical one, we inspected the data looking for neurons able to respond comparably to contra- and ipsilateral row A-whisker stimulation, namely side-invariant responding neurons^6^. Notably, in the three dimensional space individuated by the raw response parameters (i.e., peak delay, amplitude and slope), the pool of ipsilateral responses tends to overlap with the contralateral one in plBC but not in amBC (Fig. 6a,b). To quantify the similarity between ipsi- and contralateral responses in the two BC subregions, parameters were normalised in the z-distribution and the Mahalanobis distance was measured for each response type. We found that contra- and ipsilateral plBC responses showed low values and concentrated around the bisector representing equidistant points from both response types (Fig. 6c). In contrast, contra- and ipsilateral amBC responses were clearly separated (Fig. 6d), confirming that the two responses were remarkably similar in plBC but not in amBC. Accordingly, the distance of ipsi-from contralateral excitatory responses (Fig. 6e) was much shorter in plBC than amBC neurons (p=1.2·10^-6^). An analogous result was obtained for inhibitory responses (Fig. 6e) when pooled contralateral (n= 7) and separated ipsilateral responses from plBC and amBC were compared (p=9.7·10^-4^), extending this contra-/ipsilateral similarity up to the pattern of inhibition. We next computed the cross-correlation coefficient between contra- and ipsilateral excitatory responses for each neuron (Fig. 6f). Correlation was significantly higher for plBC than amBC neurons (p=1.3·10^-7^). Moreover, both measures changed dramatically when comparing responses belonging to plBC neurons between CTRL and TTX (Fig. 6e,f), confirming a role for the CC in producing the similarity between the two responses. Lastly, in more than one third of plBC neurons (34%, 15/44), the cross-correlation between contra- and ipsilateral responses exceeded 0.95, while in amBC it was only 3.8% (1/26). Such plBC neurons were unevenly distributed along the cortical depth, with the majority concentrated in L5/6 (Fig. 6h) and showing an ipsilateral response indistinguishable from the contralateral one (Fig. 6g). Notably, their rate and probability of action potential firing were also statistically equivalent, in contrast with remaining plBC neurons that were more sensitive to the contralateral side (Fig. 6i,j). These data demonstrate the presence of row A side-invariant responding neurons restricted to the plBC and suggest their involvement in the representation of the midline of the whiskers’ sensory space.

**Fig. 6.**
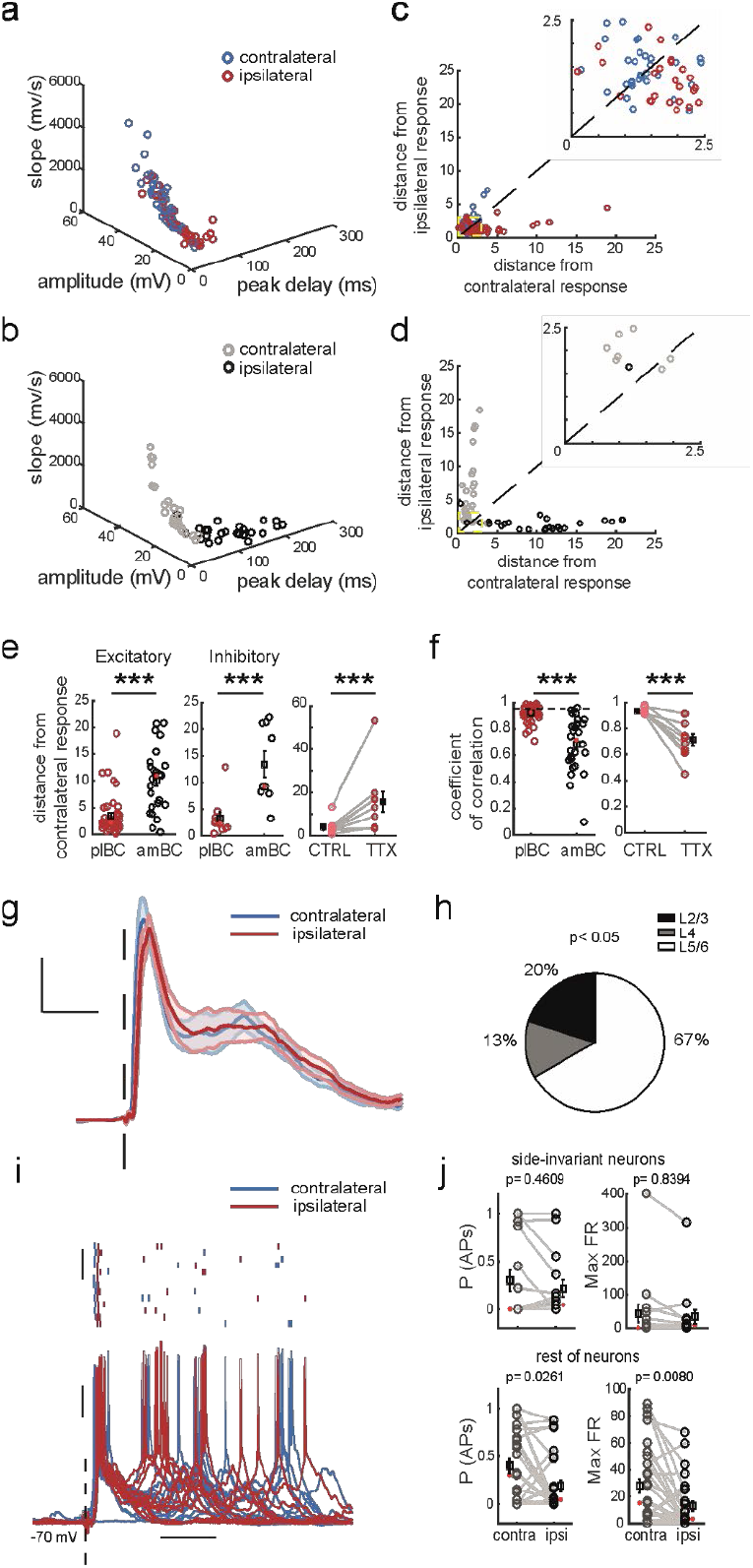
Side-invariant representation of row A-whiskers. **(a)** Parameters of contra- and ipsilateral plBC responses to row A-whiskers stimulation. **(b)** Parameters of contra- and ipsilateral amBC responses to row E-whiskers stimulation. **(c)** Mahalanobis distance (a.u.) in plBC. Inset shows magnification within 2.5 a.u.. Colour-code as a. **(d)** Same as c in amBC. Colour-code as b. **(e)** Statistical comparisons of ipsi-to-contralateral distance (x-axis in c and d) for the excitatory (left) and inhibitory (centre) response components, plus the effect of TTX on the excitatory one (right) [exc. distance (a.u.): plBC=2.2 (1.6-3.7), amBC=11.0 (5.4-12.8), p=1.2·10^-6^; inh. distance (a.u.): plBC=2.3(1.4-3.4), amBC=9.2(8.1-21.1), p=9.7·10^-4^; WSR test on exc._TTX_ distance (a.u.): CTRL=2.7(2.4-3.5), TTX=14.8(3.9-17.0), p=0.0020]. **(f)** Cross-correlation coefficients between contra- and ipsilateral waveform averages of single neurons (left) and the effect of TTX (right) [coeff._contra/ipsi_: plBC= 0.92 (0.88-0.96), amBC= 0.71 (0.56-0.83), p=1.3·10^-7^; WSR test on coeff._contra/ipsi_ in TTX: CTRL=0.92 (0.90-0.96), TTX=0.70 (0.62-0.83), p=0.0020]. Horizontal dashed line indicates 0.95. **(g)** Grand average ± S.E.M. (shadows) of contra- and ipsilateral row A-whiskers responses aligned at stimulus delivery time (dashed line) with correlation in f >0.95. Vertical bar= 5 mV, horizontal bar= 100 ms. **(h)** Side-invariant responding neurons in g preferentially occupy L5/6 [df=2, *χ*^2^ =7.60, p<0.05]. **(i)** Example raw trace of a side-invariant spike-responding neuron (bottom) with rasterplot (top). Responses are aligned at stimulus delivery time (dashed line). Vertical bars= 5 trials (top), 10 mV (bottom); horizontal bar= 100 ms. **(j)** Quantification of firing activity following row A-whiskers stimulations.

### The posterolateral region receives heterotopic contralateral synaptic inputs more robustly than the anteromedial one

While showing a preference for a given whisker, BC neurons can also respond to the stimulation of whiskers other than the preferred one^46,47^. Critically, integrating whisker inputs across the whisker array allows BC neurons to represent complex features of objects in the somatosensory space^48,49^. In order to represent in unison the right and left whiskers’ sensory hemi-spaces, such an operation may be required across the hemispheres^7^. The presence of callosal heterotopic connectivity would allow the interaction between non-preferred whisker representations of the two BCs. In order to explore this possibility, we analysed the heterotopic projections of plBC and amBC after BDA injections. Our result showed a scant presence of heterotopic axons reaching the opposite hemisphere at non-mirror symmetric regions, respectively in amBC and plBC (Fig. 7a,b), with a tendency to be slightly higher in plBC (Fig. 7c, p=0.073). Next, we examined synaptic responses of callosal heterotopic connectivity. We expressed ChR2 in the BC of NEX-Cre mice and compared the evoked responses to the homotopic, heterotopic, or both simultaneously (BC_wide_) light activation of the contralateral BC (Fig. 7d–k). In addition to the recorded neurons, we also obtained extracellular LFP recordings in the stimulated BC to ensure the optogenetic activation pattern (Fig. 7f, g). Notably, while 100% (4/4) of plBC neurons could respond to BC_he_-stimulation, a response was detected only in 57% (4/7) of amBC neurons, suggesting that plBC neurons are more functionally connected with heterotopic contralateral territories than amBC ones. Indeed, in amBC, BC_he_-stimulation evoked responses at longer onset delays than the BC_ho_-stimulation (p=0.0084; Fig. 7h,i & Sup. Tab. 5). By contrast, onset delays did not differ in plBC (p=0.2637; Fig. 7j,k). The high proportion of responding neurons and the absence of onset delays differences suggest that heterotopic CC-fibres can activate plBC neurons as directly as the homotopic ones, while this activation is rarer and likely mediated by a more indirect pathway in amBC neurons. Nonetheless, responses to BC_he_-stimulation were generally weaker than the BC_ho_- and BC_wide_-stimulation in plBC neurons (Fig. 7j,k): amplitude and slope were reduced, and the maximal peak was delayed in this condition (Sup. Tab. 6). Together, the anatomical and electrophysiological data show that the heterotopic innervation is weaker than homotopic innervation in plBC. Lastly, responses to BC_wide_-stimulation were slightly more vigorous than BC_ho_-stimulations (Fig. 7h–k), suggesting a certain degree of inputs summation between homo- and heterotopic contralateral territories (Fig. 7j,k). Yet responses did not differ statistically, confirming that the two BCs mainly operate in a homotopic fashion, with a small heterotopic contribution.

**Fig. 7.**
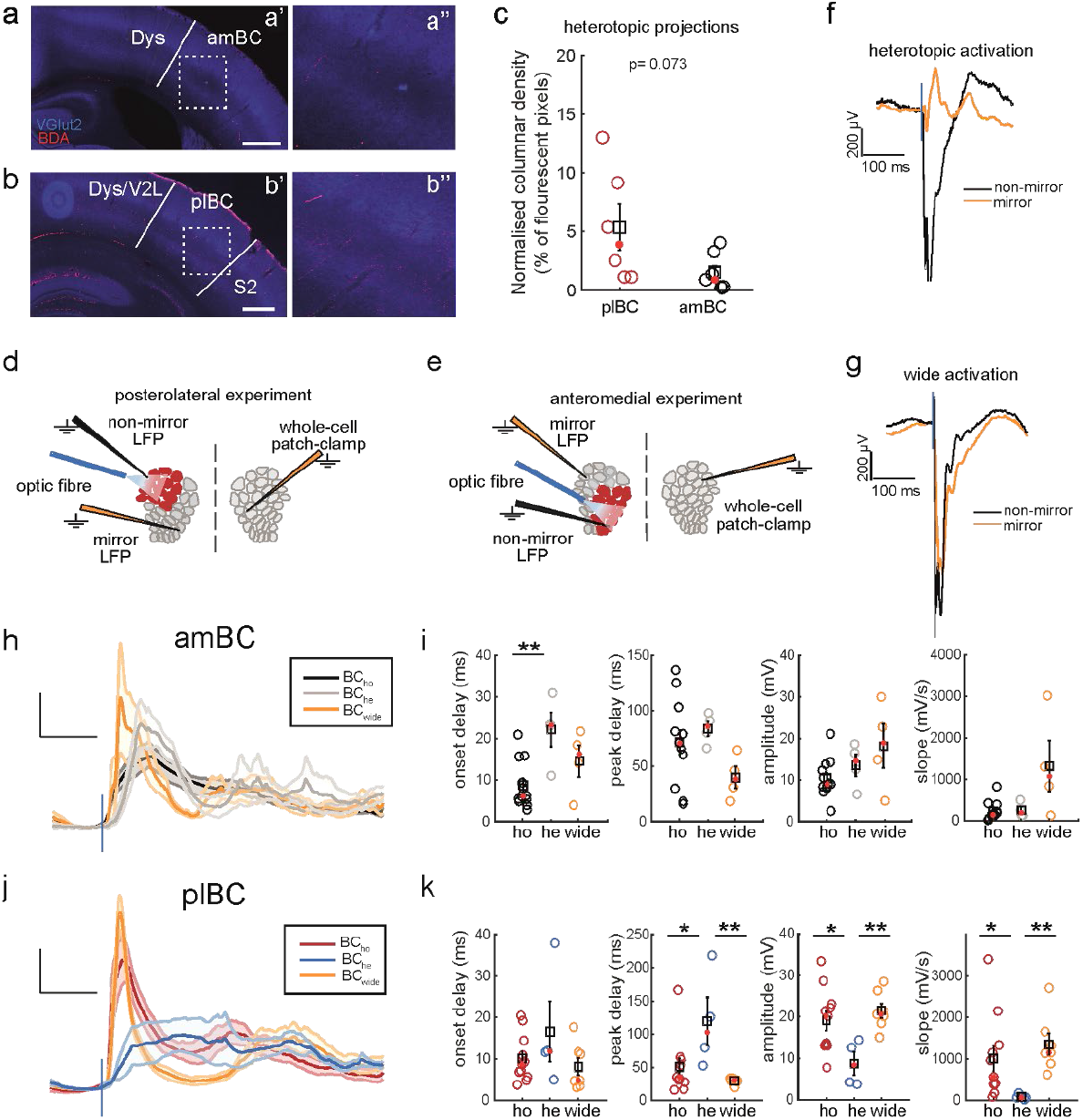
Homotopic, heterotopic and wide callosal activation of the barrel cortex subregions. **(a)** Heterotopic axons in amBC from contralateral plBC BDA injection (a’) with inset from dashed square (a’’). **(b)** Heterotopic axons in plBC from contralateral amBC BDA injection (b’) with inset from dashed square (b’’). Note that contralateral BDA injections are from the same brains of Fig. 1a,b. **(c)** Statistical comparison of heterotopic axonal fluorescence in amBC and plBC. **(d)** Schema of the experimental setting for heterotopic optogenetics stimulation and recordings in plBC. **(e)** Same as d for amBC. **(f)** Example waveform averages of mirror and non-mirror LFPs in heterotopic experiments aligned at stimulus delivery time (blue line). **(g)** Same as f for wide activations of the BC. **(h)** Grand average ± S.E.M. (shadows) of optogenetics responses aligned at stimulus delivery time (vertical blue line) in the three conditions in amBC. Note that homotopic response is the same shown in Fig. 5g. **(i)** Statistical comparison of parameters for responses in h [KW onset delay (ms): df=2, *χ*^2^=7.09, p=0.0084]. Fisher’s LSD post-hoc test values for i and k are provided in Sup. Tab. 5. **(j)** Same as h in plBC. **(k)** Statistical comparison of parameters for responses in j [KW onset delay (ms): df=2, *χ*^2^=2.67, p=0.2637]. Scale bar a’-b’= 500 μm.

## Discussion

Studies on the callosal contribution to BC activity^11–16,32^ have so far disregarded how ipsilateral responses relate to the callosal innervation arrangement^19–25^. To our knowledge, the only study in which whiskers of different rows were stimulated and ipsilateral responses were recorded did not include the stimulation of row A^14^. Available data on this circuit is provided by ref.^40^, where *ex vivo* monosynaptic callosal connections were mapped throughout the cortical depth in the row A representation. Yet, such a preparation did not allow recording of the associated sensory responses. Here we show that the row A representation in plBC is the most active despite the block of thalamocortical inputs during mouse exploratory behaviour. Bilateral trimming of the whiskers was not enough to refrain the BC thalamocortical activation (Fig. 1e). Curiously, the anteroposterior extent of c-Fos expression differed between intact whiskers (A< P) and trimmed whiskers (A> P) cohorts (Sup. Fig. 1a,b). Unlike in the posterior aspect, in the anterior aspect of the BC, the representation of the furry skin intervening between whisker follicles overlaps with that of small, anterior whiskers^50^. Together with our common observation that mice with bilaterally trimmed whiskers touched cage walls and littermates directly with the snout, these results could reflect a different exploration strategy. We hypothesise that, while mice with intact whiskers mostly used long whiskers during exploration, mice with trimmed whiskers used the common fur on the anterior snout, thereby activating the barrels representing short whiskers. Such an activation is not reported normally for mice left with only one whisker. Perhaps, the spared whisker allows mice to maintain a proper distance between snout and surfaces, an otherwise difficult task. Further behavioural studies are needed to confirm this interpretation. Nevertheless, lidocaine injection in the whisker-pads could prevent such activation and be used to isolate the ipsilateral contribution to the activity of BC subregions, which remarkably overlapped with the pattern of callosal innervation (Fig. 1a,d–g, 5a).

### Neural pathways

Contralateral stimulations activated L4 faster than L2/3 and L5/6. This suggests the recruitment of the lemniscal pathway involving the dorsomedial part of the ventroposteromedial thalamic nucleus. By contrast, deeper layers tended to respond faster than superficial layers following ipsilateral stimulations. Such a trend was stronger in plBC, whose L5/6 neurons responded earlier than in amBC (Fig. 2g). This may result from the higher abundance of callosal innervation reaching the plBC that can increase the likelihood of monosynaptic responses in the recorded neuronal pool, thereby shortening the average response onset. Indeed, the same tendency to show a faster response onset in plBC than in amBC is present for L2/3 (p=0.0545, *d*= 0.92, MD= 13.7 ms), which also receives monosynaptic input from contralateral L2/3 in the representation of row A^40^. Regarding L4, barrel hollows are virtually devoid of functional CC contacts, at least from axons originating in L2/3 of the opposite hemisphere^40^, while agranular septal compartments are often reported to be the recipient of CC axons in this layer^17,22,23,51,52^. Hence, a population trend towards indirect, multisynaptic responses can explain why plBC and amBC responses in L4 have more similar onset delays (Fig. 2g). A stronger input to L5 than to L2/3 of the row A representation characterises callosal axons originating in L2/3 of the opposite hemisphere^45^. This can explain the concentration of side-invariant responding neurons in the L5/6 pool (Fig. 6h) and their increased tendency to fire upon ipsilateral stimulation (15/19, ~80%). In this vein, the increased amplitude of thalamocortical responses in plBC of callosally-projecting L2/3 (Fig. 2d,e & Sup. Fig. 1b,c) may also contribute to the vigour exhibited by L5/6 ipsilateral responses. Instead, the weaker ipsilateral response of amBC neurons (Fig 2f,g) may result from fewer direct callosal connections or from a reduced myelination/calibre of the callosal axons entering this region (see below). In the former case, widespread subthreshold responses may result from inputs travelling *via* local axon collaterals of neurons monosynaptically connected to the CC, as previously proposed^3^. Since most of the CC afferents in the BC enter septal agranular areas, monosynaptically connected septal neurons would be the best candidate for this mediation in BC areas with scant CC innervation.

### Interhemispheric transmission

Ipsilateral responses produce fewer spikes than contralateral ones in S1^14–16^. Consistent with this fact, in previous^11,12^ and in the present work, most subthreshold ipsilateral responses rose slowly and with a smaller amplitude than their contralateral counterparts, leading to a minor drive for firing. Callosal conduction velocities show great variability^53^ due to the rich diversity in axonal calibre and myelination^2,3^. Moreover, in the mouse S1, infragranular layers are more extensively myelinated than supragranular ones^54^, with both compartments containing callosally projecting neurons^17^. Such heterogeneity may provoke an asynchronous synaptic summation of inputs in postsynaptic neurons translating into a minor drive for spiking. Indeed, without even considering L5/6, the stimulation of more than one callosal axon belonging to L2/3 neurons led to such an asynchronous input summation in postsynaptic row A neurons^40^. This break of input synchrony through the CC, more evident in scantily connected areas, reduces the strength of the original thalamocortical activation of the opposite hemisphere. Such synchrony disruption could be stronger during whisking, as suggested by a previous description of activity desynchronisation between BCs that were recorded at BC coordinates roughly corresponding to our amBC ones^55^. This weaker input may instruct brain areas which integrate the S1 output about the side of a contacted object in the somatosensory space. For example, comparably to the BC, the sensorimotor dorsolateral striatum shows weaker whisker responses to ipsi-than to contralateral stimulation^12^. By contrast, increasing the strength of callosal connectivity may mitigate this phenomenon such that thalamocortical activations can be replicated in the opposite hemisphere. In the representation of midline body parts, this neural operation is thought to maintain the continuity between right and left sensory maps^6,9,10^. Here we propose such a scenario for row A-whiskers, which show a significant degree of side-invariant representation in the BC (Fig. 6a, c, f–g), regarding excitatory and inhibitory components of the subthreshold whisker responses (Fig. 6e), evoked spiking activity (Fig 6i–l), and the highest degree of callosal innervation spared from developmental pruning^24^.

### Beyond the midline rule in the whiskers system

The side-invariant representation of row A-whiskers suggests that they have less discriminative capabilities than remainder whiskers. This is because spatial coding, which relies on the topography of single whiskers within the array^56^, may be less efficient in discriminating which one of the two rows A contacts an object. This accompanies a difference in muscles’ anchoring to row A-follicles, which lead them to counterrotate during torsion in the whisker-cycle with respect to remainder whiskers^57,58^, suggesting a unique role for these rows. Here we propose two possible scenarios. On the one hand, the side-invariant representation of row A-whiskers may allow the animal to gather positional cues of tactile features belonging to an object in the scanned space with reference to the facial midline. This is mainly suggested by the organisation of the neural pathways. Indeed, such maps of the space are most prominently represented by L2/3 and L5 neurons in the BC^48^, with both compartments projecting/receiving axons trancallosally^17^. Moreover, plBC at the S1/S2 border receives a more direct heterotopic input than amBC (Fig. 7h, i) in addition to the homotopic input, thus potentially constituting the earliest connectional hub for the two sensory hemi-spaces. Finally, the S1/S2 border sends ipsi- and contralateral projections to the perirhinal cortex^25^, a polymodal area involved in object recognition^59^, where an egocentric map of objects’ features may be built up in association with other modalities. In this manner, the animal could explore objects through the position of their features with respect to its face. A second use of this side-invariant representation of row A-whiskers may be for the easy transformation of frame of coordinates between body parts lying on the facial midline. These transformations are needed when one system has to approach the same object in the space that another system has detected^56^. For example, the rows A may allow the animal to accurately position its nose over a tactually salient spot in space. This second interpretation is supported by the exemplary case of the star-nosed mole, whose life heavily relies on palpation. To align the mouth with food, the star-nosed mole performs a ‘tactile foveation’, which is mediated by extremely touch sensitive, midline appendages interposed between the two nostrils^61^. Strikingly, by resembling the row A-whiskers representation in mice and rats, the midline appendages’ S1 representation has the densest callosal innervation among the peers composing the tactile ‘star’^62^. Hence, the similarity in sensory modality, midline positioning and callosal representation may result in an analogous behavioural role in these species.

## Methods

### Ethical approval

All the experimental procedures were in conformity with the directive 2010/63/EU of the European Parliament and of the Council, and the RD 53/2013 Spanish regulation on the protection of animals use for scientific purposes, approved by the government of the Autonomous Community of Valencia, under the supervision of the Consejo Superior de Investigaciones Científicas and the Miguel Hernandez University Committee for Animal use in Laboratory.

### Experimental animals

For c-Fos experiments, we used wild-type C57BL6 mice (n= 16) of either sex between 2 and 3 months of age. For electrophysiological experiments we used wild-type C57BL6 mice (n= 23), and the transgenic line NEX-Cre (Goebbels et al., 2006) (n= 23) mice (generously donated from Dr. Klaus-Armin Nave and Dr. Victor Borrell) of either sex and aged 2-6 months. All animals were housed preferentially with cage-mates at T= 24° C in our institution’s animal facility with 12h daily illumination and food and water *ad libitum*.

### c-Fos immunoassay

In order to habituate mice to the environment, the first day they were placed in a covered empty arena (black PVC material; 50 x 50 x 70 cm) in complete darkness for ~1 hour. The following day, they were briefly (~15 min) anaesthetised with 2-4% isoflurane in oxygen (0.8 L/min) to allow interventions on whiskers pads. In trimmed whiskers condition, whiskers were trimmed bilaterally, paying attention to reach the very base of the whiskers. For bi- and unilateral lidocaine conditions, lidocaine (150-200 μL) was injected subcutaneously, with the volume subdivided in one to two injections in order to cover the entire whisker pad. We waited 90-120 min for the anaesthesia to fade completely before testing the mice in the arena in the presence of new objects of various sizes and shapes. Success of lidocaine injections was assessed by visually checking the inability to whisk in the injected side(s) prior to test. In the majority of experiments, animals were tracked with a Pixy camera to monitor the locomotor performance (Sup. Fig. 1c). After 45 min of free exploration, mice were sacrificed by lethal injection of sodium Pentobarbital, transcardially perfused with 4% paraformaldehyde in 0.1 M phosphate buffer (pH 7.4), decapitated and their brain was extracted. After tangential sectioning and c-Fos immunostaining of flattened cortical preparations, images were acquired with a confocal microscope (Leica SPEII) with 10 μm step-size using a 20x immersion objective. Data from images were quantified using the Fiji software. Area and disposition of ROIs from barrels and septa were recognised by VGlut2 immunostaining, extracted and then projected onto the respective image showing the c-Fos channel. In this manner, we could assign the count of immunopositive nuclei to their BC compartments. By dividing the nuclei count for the ROIs area, we obtained a measure of c-Fos^+^ nuclei density (#nuclei/ 0.1 mm^2^) in target barrels (A_1_-A_4_, B_1_-B_3_, C_1_-C_7_, D_1_-D_7_, E_1_-E_7_) and septa (A/B, B/C, C/D, D/E). Anterior BC contained A_4_, B_3_, C_4_-C_7_, D_4_-D_7_, E_4_-E_7_ barrels, while posterior BC contained A_1_-A_3_, B_1_-B_2_, C_1_-C_3_, D_1_-D_3_, E_1_-E_3_ barrels. In lidocaine cohorts, if the contralateral BC showed c-Fos expression clearly resembling the one of thalamocortical activation (i.e., very dense c-Fos labelling within the barrel hollow) in the barrels of interest, the lidocaine effect was considered faded prior to sacrifice and mice were excluded from the analysis. Count in missing barrels of histological sections was replaced by NaN.

### Viral injections

To impose the activation of callosal-projection neurons, we infected pyramidal cells of the right BC with AAV viral vectors encoding optogenetic tools. Anaesthesia (2-4% isoflurane in oxygen, 0.8 L/min) was induced and maintained for the course of the surgery in NEX-Cre mice immobilised in a stereotaxic apparatus (Kopf Instrument). Lidocaine cream was applied to the skin over the skull prior to opening and eyes were covered with ophthalmic gel (Viscotears 2mg/g) to prevent corneal desiccation. Analgesic (Metacam, 0.4 mL) was delivered by intraperitoneal injection. Craniotomy was drilled either at coordinates AP -2 ML -3.8 (plBC) or AP -1 ML -3.2 (amBC) from bregma, exposing the dura mater. The ssAAV-5/2-hEF1α-dlox-hChR2(H134R)_EYFP(rev)-dlox-WPRE-hGHp(A) viral construct (ETH Zurich) was injected in small volumes (100-200 nL) at 0.7 mm depth with a precision injector (Nanoliter, WPI). Skin was then closed with surgical glue and animals rested in a recovery chamber until restoration of normal locomotor activity.

### Immunohistochemistry

After every experiment, mice were perfused with 4% PFA. Their brains were post-fixed also in 4% PFA for 2h at room temperature and then stored at 4 °C in PBS with 30% sucrose plus 0.1% sodium azide before their use. For flattened preparations of the cortex, the entire cortical mantle was isolated from the brain and pressed overnight in a custom-made, plastic press-and-hold apparatus of 1 mm depth in a 4% PFA bath prior to sectioning. Tangential slices 80 μm thick from flattened cortex and coronal slices 50 μm thick from intact brains were cut with a digital cryotome (Microm HM450; ThermoFisher). After washing with PBS, sections were exposed to a blocking solution of 0.1M PBS with 1% BSA and 0.5% Tryton-X 100 for 2h at room temperature. Next, they were incubated overnight with primary antibodies (4 °C) diluted in blocking solution. To stain thalamocortical afferents for recognising L4 barrels, we used guinea pig anti-VGlut2 (1:4000; Synaptic Systems). To detect c-Fos expressing nuclei, we used rabbit anti-cFos (1:700; Synaptic Systems). Mouse anti-NeuN (1:250; Sigma Aldrich) was used for neural somas membranes. The following day, sections were washed and incubated for 2h at room temperature in blocking solution with secondary antibodies, adding streptavidin for tissues previously used in electrophysiological experiments (see below). Once this time elapsed, they were mounted on microscope slides (Menzel-Gläser Superfrost Plus; ThermoFisher), covered with mowiol (Calbiochem) and coverslipped (Menzel-Gläser; ThermoFisher). To recover biocytin-filled neurons from electrophysiological experiments, we used Cy2- or Cy3-conjugated streptavidin (Jackson ImmunoResearch Laboratories), diluted, respectively, 1:500 and 1:1000 in blocking solution. As secondary antibodies, we used goat anti-guinea pig Alexa fluor 568 and 633 (ThermoFisher), both with 1:500 dilutions, and donkey anti-rabbit Alexa fluor 488 (ThermoFisher) at dilution 1:500, and DAPI (ThermoFisher) for nuclei. For cell recovery in amBC recordings, coronal sections were collected from bregma AP -0.58 to AP -1.58, while for plBC from bregma AP -1.82 to AP -2.30 (following Paxinos and Franklin, 2001).

### Axonal tracing and quantification

To label callosal projections in left barrel cortex subregions we injected 200nL of 10% biotinylated dextran amide (BDA) in the right hemisphere either in the plBC (AP −2, ML 3.9) or the amBC (AP −1, ML 2.8) subregion. After one week, animals were perfused with 4% PFA, and brains were stored at 4°C in 4% PFA overnight. As above, 50μm thick coronal cryosections were incubated overnight with pig anti-VGlut2 (1:4000; SynapticSystems), followed the second day by the application of goat anti-guinea pig Alexa fluor 633 (1:500; ThermoFisher) and Cy2-conjugated streptavidin (1:500, Jackson Immuno Research). Pictures were taken with the Thunder microscope (dry, 10x; Leica Microsystems). For each brain, seven pictures were taken: one image of the section with the highest fluorescence at the injection site, three consecutive sections containing the row E barrels (amBC), and three consecutive sections containing the row A barrels (plBC). Two different ROIs were drawn with ImageJ: one comprising the whole column of the rows A/B whiskers (AP -2; ML 3.5-4) and another comprising the whole column of the rows D/E (AP -1; ML 2.75-3.25). Images were then binarised with Otsun’s threshold and in each ROI the number of pixels corresponding to BDA fluorescence was quantified. We estimated the callosal fibres’ density by the number of fluorescent pixels divided by the total number of pixels of the ROI. Normalisation of the data was performed by dividing each density value by the average density of the injection ROI.

### Whiskers and optogenetic stimulation

With the exception of the pairs A_2_-A_3_ and E_2_-E_3_, whiskers were trimmed bilaterally at their base to avoid any unintended stimulation accompanying the one of target whiskers. Spared whiskers, separately for each row (i.e., A and E), were glued together with super glue along their length. Then, they were attached to the tip of Teflon-coated stainless wire of a custom-made solenoid puller (see Sup. Fig. 3a) with super glue. In this manner, and depending on the target cortical area (i.e., plBC or amBC), mirror whiskers of both sides of the snout were connected to the solenoid puller. Whiskers’ displacement consisted of fast caudo-rostral pulling (~3.5 mm) lasting 15 ms, delivered every 3 or 5 s, contra- and ipsilateral to the patch-clamp recording electrode. For optogenetics experiments in NEX-Cre mice, single step-like photo-stimulations lasting 5 or 10 ms were delivered every 3 or 5 seconds, with an optic fibre (400 μm diameter) connected to a blue light source (Prizmatix LED, 453 nm wavelength, 3.76 mW light power) and governed by the CED1401 (Power3) through a custom-made program (Spike2). Two craniotomies were drilled. The fibre tip was positioned perpendicular to the surface of the brain within the craniotomy used for delivering the viral construct, either at plBC or amBC coordinates. A micro-holding device^62^ was used to keep in the same coordinate the borosilicate capillary (1B150F-4, WPI) containing the LFP electrode and the optic fibre, except that the tip of the electrode was deepened 1 mm into the cortex, while the optic fibre rested on the brain surface. In the second, fibre-free craniotomy, another LFP electrode (0.9-1.2 MΩ) was placed at 1 mm depth into the cortex to check for the anatomical specificity of the optogenetics activation. Mirror- and non-mirror LFPs were considered jointly active in BC_wide_-stimulations if their negative deflection in response to optogenetic illumination surpassed 400μV (Fig. 7g).

### TTX experiments

The disposition of plBC and amBC LFPs was similar to the optogenetics settings, except that the fibre inserted in the micro-holder was replaced by a borosilicate capillary with broken tip, containing 1-1.5 μL of TTX 100 μM. TTX was released through positive pressure at 1 mm cortical depth in plBC. Responses to ipsilateral row A-whiskers stimulation were analysed after that, from plBC, the depression of activity reached the electrode in amBC (Fig. 4a).

### Electrophysiological recordings

Electrophysiological recordings were performed on top of an air table for vibration cancellation (CleanBench, TMC). Mice were head-fixed in a customised stereotaxic apparatus with ear-bars (Stoelting). We induced anaesthesia by i.p. injection of ketamine (75 mg/kg) and medetomidine (1 mg/kg) diluted in 0.9% NaCl saline solution. One third of the original dose was injected intramuscularly to maintain anaesthesia once paw reflex could be evoked or the LFP trace exhibited signatures of awakening. To avoid mechanical instability induced by respiration, a tracheotomy was performed prior to animal immobilisation in the stereotaxic apparatus. A tube blowing oxygen-enriched air was placed ~1 cm in front of the cannula, previously inserted in the trachea. The animal rested over a heating-pad governed by a thermostat (FHC Inc.) to maintain core body temperature at 36.5 ± 0.5°C. Depending on the experiment, 2 to 3 circular craniotomies were drilled over the BCs (~5 mm diameter). Typically, they were one per hemisphere at mirror symmetric coordinates (reported in ‘Viral injections’), plus another in the right hemisphere when required, to host a second LFP at non-mirror symmetric coordinates with respect to the contralateral (left hemisphere) patch–clamp electrode. Craniotomies were constantly kept wet by application of 0.9% NaCl solution for the entire recording duration. In the craniotomy for patch-clamp, we gently removed the dura with a syringe needle bent at the tip and stopped eventual bleeding with cotton sticks. To approach the cortical representation of target whiskers within plBC or amBC craniotomies, we delivered contralateral stimulations of the whiskers group matching its cortical representation, respectively, the pair A_2_-A_3_ for plBC and E_2_-E_3_ for amBC. A borosilicate capillary filled with 0.9% NaCl solution contained a LFP electrode (0.9-1.2 MΩ). We used this to navigate the craniotomy through a digital micromanipulator (Luigs & Neumann, set at 1 μm precision) for sampling cortical responses to whiskers’ stimulations on the dura’s surface. After that the average of 10-20 LFP responses was calculated online by the software (Spike2), the electrode was moved to a new position within the craniotomy, and the process repeated (Supp. Fig. 3c). After 3-6 loci sampled, by visually comparing the averages obtained, the one with the strongest initial negative deflection was chosen to be the location for patch-clamp recordings. Especially in younger animals (PD ~60), these loci could be also recognised on the brain surface with reference to brain vessels. For patch-clamp recordings, we used an angle of ~60°. Cortical depth of the recording was collected through display reading of the micromanipulator (1μm step-size). By comparing our histological preparations with display readings and the bibliography (especially ref.^40^), we considered as L2/3 neurons recorded between 101 and 300 μm from the pia, as L4 the ones recorded between 301 and 500 μm, and L5/6 neurons below 501 μm until the white matter, at approximately 1100 μm. After anatomical recovering of biocytin-filled neurons, these coordinates associated with the penetration angle showed to be optimal for neurons recorded in plBC. However, for recordings in amBC a more perpendicular angle would have been required. In Sup. Fig. 3d we provide the details of the correction procedure for depth in amBC. Pipettes for patch-clamp recordings (7-11 MΩ), were back-filled with intracellular solution containing: 125 mM K-gluconate, 10 mM KCl, 10 mM Na-Phosphocreatine, 10 mM HEPES, 4 mM ATP-Mg and 0.3 mM GTP-Na. pH and osmolality were adjusted to ~7.4 and ~280 mOsm/L, respectively. Biocytin (0.2-0.4%, Sigma Aldrich) was then added to the intracellular solution to recover the recorded cell after every experiment (Fig. 2c). Borosilicate capillaries, both used for LFP or patch-clamp recordings, were pulled with a micropipette puller (Flaming/Brown P-1000, Sutter Instrument). Recorded signals were amplified with MultiClamp 700B (Molecular Devices), converted to digital with a CED 1401 (Power 3) acquisition board at 20 KHz, and streamed to a laptop (Windows 10) running Spike2 software (Cambridge Electronics Design). At the end of each experiment, mice were sacrificed with a lethal i.p. injection of Pentobarbital, perfused transcardially with 4% paraformaldehyde in 0.1M phosphate buffer (pH 7.4), decapitated, their brain extracted and post-fixed overnight in 4% paraformaldehyde. Thus, brains were maintained in 30% sucrose solution with sodium azide at 4 °C until the sectioning day. In this study, we recorded intracellularly a total of 110 left-BC neurons from 54 mice (~2 neurons/mouse), including histologically recovered and unrecovered neurons. Of this pool, whisker responses were obtained from 75 neurons, and optogenetics responses from 45 neurons, 10 of which also underwent sensory stimulation. Two L4 neurons have been excluded for very long response delays (> 15 ms; see ref.^46^), indicating that the recording was misplaced. Moreover, we could clearly detect only two putative fast-spiking neurons in our pool (~3%) by the remarkably short spike half-width (< 0.7 ms; population median: 1.15 ms). Yet, their proportion has to be higher since the normally reported proportion is 10-15% for the rodent neocortex^63^. The two putative fast-spiking neurons were included in the analysis.

### Data analysis

We analysed data with Matlab 2015a (Mathworks). First, we downsampled acquired signals to 10 KHz. Then, we isolated spontaneous up- and down-states of the V_m_ through a moving average window, considering up-state V_m_ values exceeding a threshold obtained by summing the mean V_m_ within the moving window to 0.4 times V_m_ standard deviation, and down-states the values below this threshold. BC up-states often display complex shapes, characterised by 2 or more peaks^64^. Thus, short hyperpolarising events (<150 ms) falling below the threshold and thus classified as brief down-states, were elided and considered voltage excursions within up-states. Analogously, too-short depolarising events (<15 ms) misclassified as up-state were included in down-state. Neural responses to whiskers and optogenetic stimulations were isolated from 200 ms before to 500 ms after the stimulation trigger. We selected responses in down-state only for stimulations falling 100 ms after the end of an up-state. Both kinds of responses were spike-filtered with a median plus the Savitzky-Golay filters (Matlab functions: *medfilt1* and *sgolayfilt*) applied in this order. For every neuron, we obtained the waveform average through the mean of the spike-filtered responses, with spikes vectors collected separately. Onset delay of responses in down-state was defined as the delay between stimulation delivery (digital trigger) and threshold crossing by the V_m_. Threshold was defined as the mean V_m_ during 50 ms of down-state preceding stimulation multiplied by 50 times its standard deviation. Peak delay was defined as the difference between stimulation delivery and the V_m_ maximal amplitude time points, within 300 ms post-stimulation. Amplitude was defined as the V_m_ difference between the peak of the spike-filtered response and the pre-stimulus baseline (i.e., down-state or resting V_m_). Slope of the rising phase was defined as the angular coefficient of the first-degree polynomial fitted to the waveform average in the time between onset and peak delay (Matlab function: *fit*). These measures were visually assessed on the waveform averages and manually adjusted when needed. Probability of action potential firing [P(AP)] was calculated in a post-stimulus time window of 50 ms for contra- and 80 ms for ipsilateral responses, by dividing the number of times that a neuron fired at least one spike following stimulation by the total number of stimulations delivered. A given neuron was classified as spike-responding if it had a P(AP)> 0. Firing rate was computed with 20 ms bins in response to each stimulation of a protocol and the maximum extracted, yielding the maximal firing rate (max FR). Amplitude of downward deflections of the LFPs (spontaneous up-states), was obtained by smoothing the signal and using *findpeaks* in Matlab, thus averaging the values for single recordings, separately for CTRL and TTX. Peak delay, amplitude and slope were z-scored to measure the Mahalanobis distance between homotopic ipsi- and contralateral responses in plBC or amBC, both for excitatory and inhibitory components (Matlab function: *mahal*). Distance values are provided as the square root of the function output (given in squared values). Cross-correlation coefficients between ipsi- and contralateral responses were calculated in the 400 ms after stimulation (Matlab function: *xcorr*), limiting the lag range to ±50 ms. Membrane properties were calculated from negative and positive injection currents (i.e., from -100 to 100 nA, by 14 increasingly positive steps) delivered to neurons during spontaneous activity, separately for up- and down-states. Membrane constant *τ* (ms) was defined as the time at which the membrane potential reached the 37% of its resting value upon current injection, averaged through negative and positive current steps. We obtained the resistance (MΩ) through the angular coefficient of the first-degree polynomial fitted to V_m_ values receiving the current injections, and separately calculated the values for up-(R_up_) and down-states (R_down_). Finally, we measured the cells’ capacitance (pF) by dividing *τ* for the resistance in up-(C_up_) and down-states (C_down_).

### Statistics

Graphs, quantification and statistical tests have been performed in Matlab R2015a (Mathworks). Values are expressed as median (25^th^-75^th^ quantiles), with the median graphically represented by a red dot and the mean by a black square ± S.E.M. (whiskers). We used a two-sided Wilcoxon rank sum test for statistical comparison of unmatched samples and two-sided Wilcoxon sign rank test (WSR) for matched samples. To test the homogeneity of distributions we used the *χ*^2^ –test and the tabulated values to infer statistical significance, applying the Yate’s correction for df=1. Effect size was expressed as mean difference in standard deviation units, indicated as *d*. MD indicates difference between medians. For comparisons involving more than two sample distributions, we used Kruskal-Wallis (KW) test with Fisher’s Least Square Difference (LSD) post-hoc test to infer statistical significance of pairwise comparisons. We applied the Bonferroni-Holm correction in multiple comparisons. For all statistical tests, the null hypothesis was rejected at the 5% significance level. Symbols: * indicates p<0.05, ** indicates p<0.01 and *** indicates p<0.001.

## Acknowledgments

We are grateful to Dr. Klaus-Armin Nave and Dr. Victor Borrell for the transgenic line NEX-Cre (Goebbels et al., 2006), Dr. N. Anton-Bolaños for initiating R.M. to the immunohistochemistry of the barrel cortex. We thank V. Rodriguez-Milan of the Instituto de Neurociencias workshop for helping with the construction of the whiskers stimulator. We are also grateful to J. Esteve-Agraz for his help with the design of some figures. Finally, we want to thank Dr. Emilio Geijo, Dr. Juan Perez-Fernandez, Dr. Isabel Pérez-Otaño and Dr. Kevin Caref for critical reading, discussions and suggestions on this article. R.M. is supported by la Caixa-Severo Ochoa programme [2016/00006/001]; A.A-A. is supported by the ACIF-GVA programme; J.A-C is supported by the CSIC-Severo Ochoa Excellence Programme of the Instituto de Neurociencias [SEV2013-0317]; The study was supported by MINECO [PGC2018-094457-B-I00] and CSIC-Severo Ochoa Excellence Programmes of the Instituto de Neurociencias [SEV-2013-0317 and SEV-2017-0723].

## SUPPLEMENTARY INFORMATION: FIGURES AND LEGENDS

**Supplementary Fig. 1.**
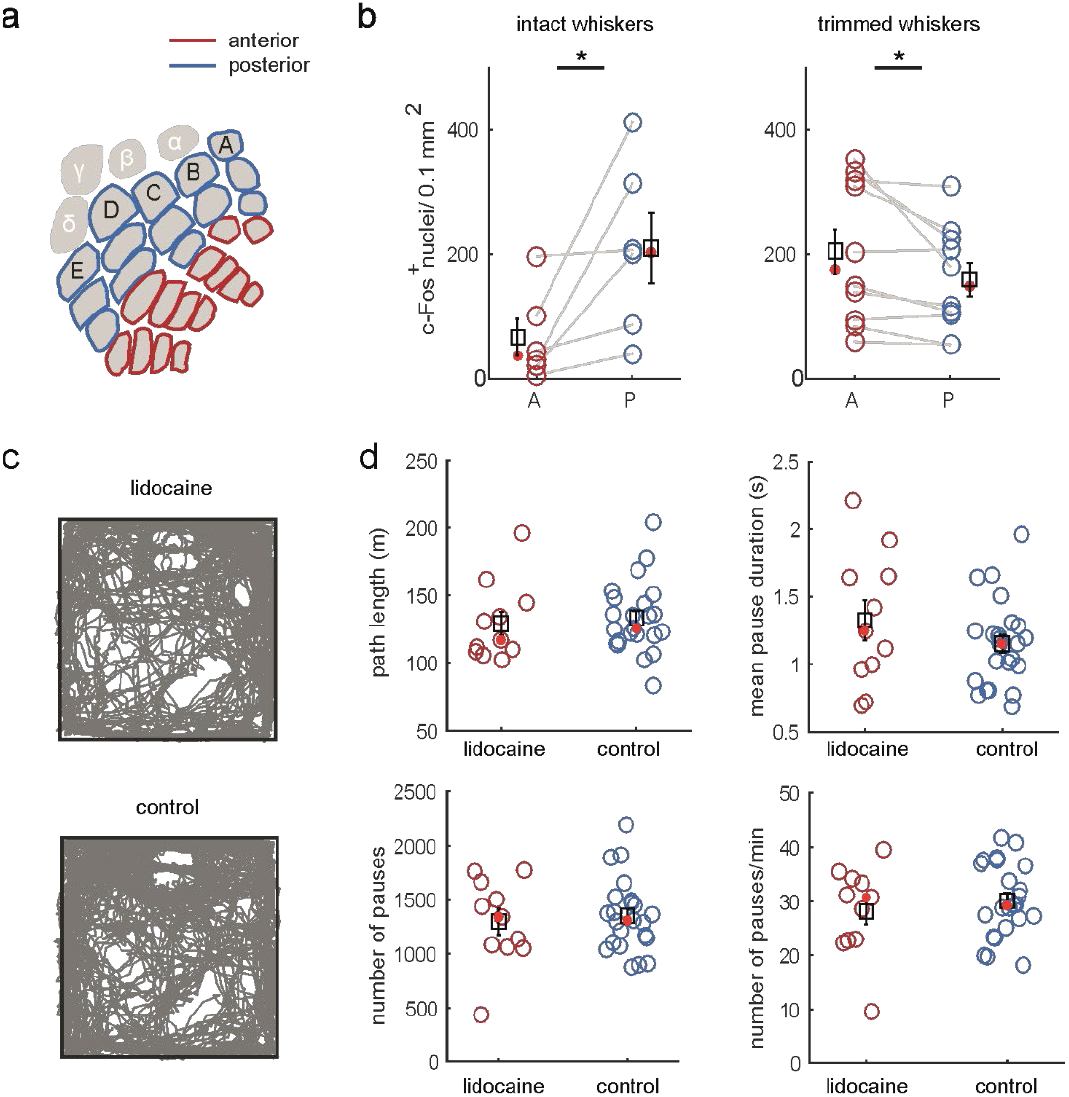
Comparison of barrel cortex c-Fos density between intact and trimmed whiskers mice cohorts. **(a)** Outline of the BC subdivision used for the definitions of anterior and posterior barrels (see Methods). **(b)** Intact whiskers mice cohort shows higher c-Fos expression in the somatotopic representation of posterior rather than anterior whiskers [WSR density: A=38.1 (20.9-102.0); P=204.0 (86.9-314.6); p=0.0313], while trimmed whiskers mice cohort shows lower c-Fos density in the somatotopic representation of posterior rather than anterior whiskers [WSR density: A=176.7 (95.0-319.8); P=149.0 (102.4-224.9); p=0.0137]. **(c)** Example tracked paths from a lidocaine-receiving mouse (lidocaine) and from a lidocaine-free mouse (control). In c-Fos and in extra pilot experiments, mice (11 lidocaine-recipient vs 23 lidocaine-free) were tracked during novel object exploration to assess the effect of lidocaine on the exploratory drive. Body position was tracked through a Pixy camera (sampling freq.= 5 Hz; pixycam.com), with the *x* and *y* coordinates streamed to an Arduino and collected with a custom-made Matlab script running on a laptop. The arena was illuminated with IR light, with the mouse carrying an IR reflective sticker glued on the top of the back to increase the signal-to-noise ratio. In lidocaine-recipient animals, lidocaine did not significantly affect the exploration drive compared to controls because no statistical difference could be detected in locomotor performance measured as length of the path, pauses count, frequency and duration. **(d)** Quantification of locomotor performance in the two conditions of c.

**Supplementary Fig. 2.**
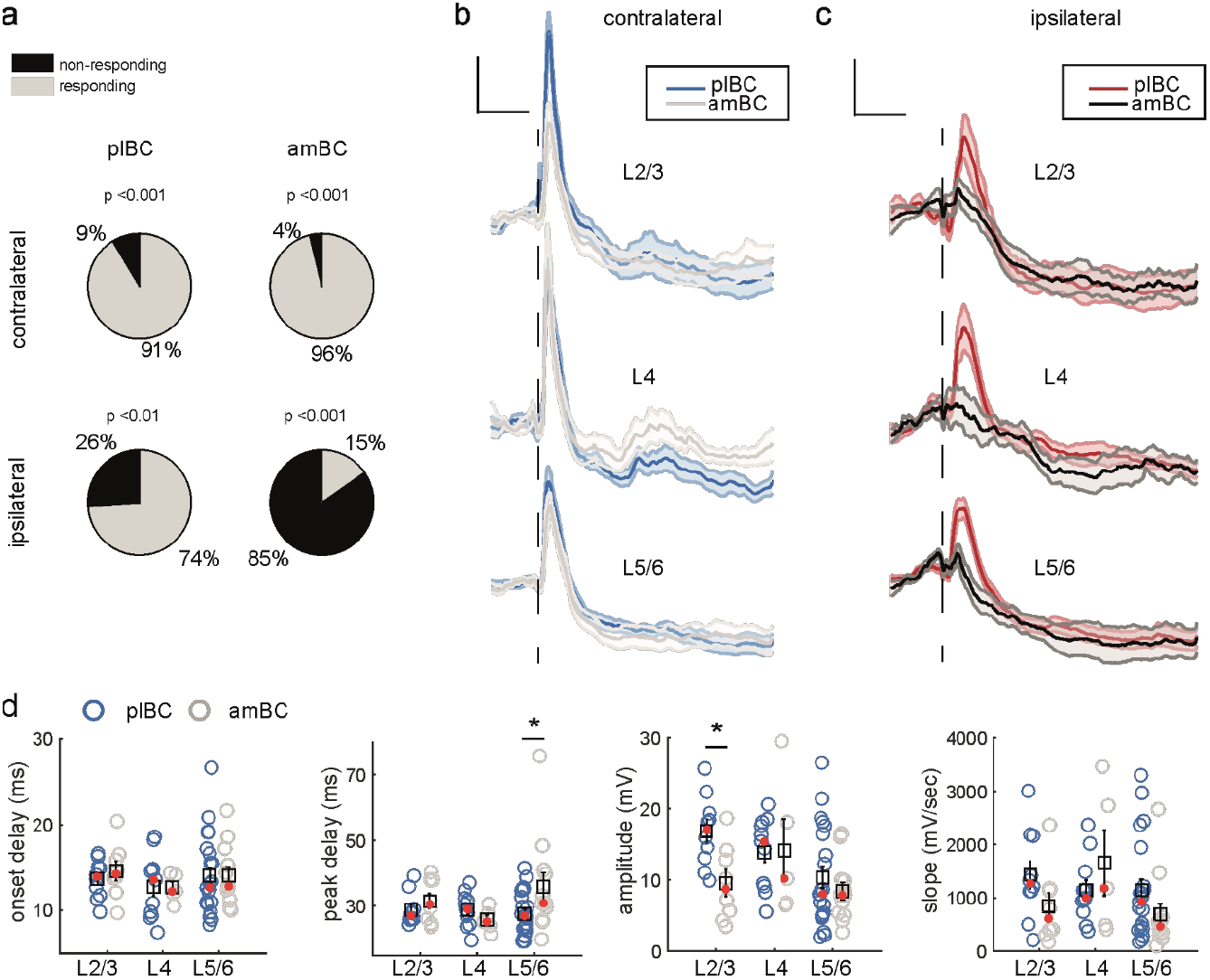
Characterisation of up-state responses. **(a)** Proportion of neurons able to respond during up-state following somatotopic ipsilateral stimulations (bottom row) is the minority in amBC (15%) but the majority in plBC (74%). P-values indicate significance at the chi-squared test for df=1. **(b)** Grand average responses ± S.E.M. (shadow) during up-state in responding neurons following somatotopic contralateral stimulation along the cortical depth, aligned at stimulus delivery time (dashed line). **(c)** Same as d for ipsilateral somatotopic stimulations but including non-responding neurons in the grand average. Note, the low number of amBC responding neurons per layer did not allow comparing parameters with plBC. Vertical bar= 5 mV, horizontal bar= 100 ms. **(d)** Statistical comparison of parameters for responses in b.

**Supplementary Fig. 3.**
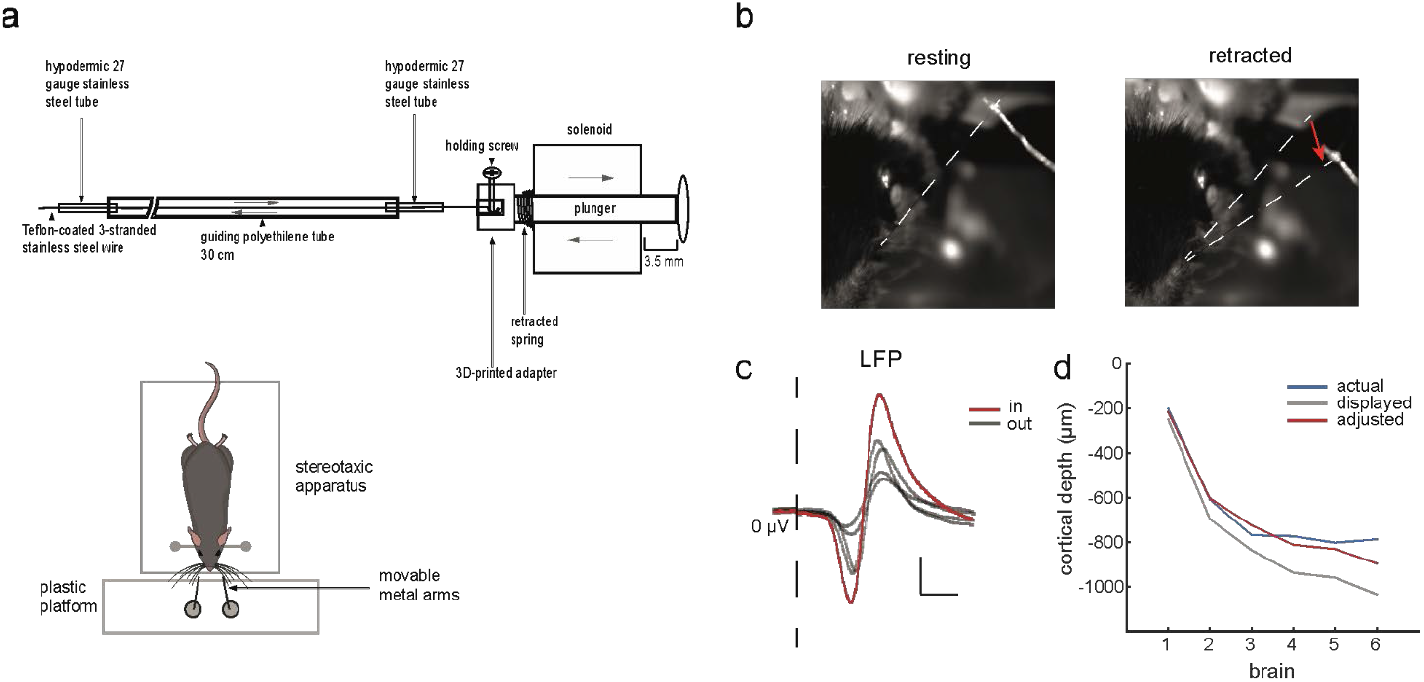
Whiskers puller, cortical mapping and neuronal depth estimation. **(a)** Whiskers’ puller. The system is composed of mechanical and electrical components governed by an Arduino. System’s assemblies at the end of a stimulation (i.e., retracted position) are shown. Whiskers’ displacement is obtained through 2 miniature solenoids (5V, 3.5 mm travel) of the pull type (ROB-11015, Electrónica Embajadores), that pull a Teflon-coated, 3-stranded, stainless steel wire (152 μm outer diameter; 3SS-2T, Science Products GmbH), glued to target whiskers at the opposite end. Such wire slides back and forth inside a guiding polyethylene tube (279 μm inner diameter, 609 μm outer diameter; 800700, Science Products GmbH). At each end of the polyethylene tube is inserted by the tip a stainless steel guide tube (191 μm inner diameter, 404 μm outer diameter; 833200, Science Products GmbH) to confer rigidity at the endings and allowing to block all the tubing complex containing the sliding wire. Miniature solenoids were mounted on a 3D-printed support platform where the stainless steel guide tube lying in front of the head of the plunger was clamped. Here, also the body of the miniature solenoid was completely blocked inside a custom-made hard-plastic pocket. Another plastic platform was mounted on the vibration cancellation table used for electrophysiological recordings, just in front of, but not in contact with, the stereotaxic apparatus hosting the mouse. From this platform, 2 metal bars emerged perpendicularly to the table’s surface, each of which carrying a movable metal arm parallel to the table’s surface and ending just in front of the animal’s snout. These arms were used to clamp the second stainless steel guiding tube next to the stimulating end of the wire and could be manually aligned and fixed in register with the whiskers. As aforementioned, the tip of a single wire was glued to the target whiskers group. Thus, electrifying the miniature solenoid provoked the retraction of the plunger that consequently pulled the whiskers group. In this manner, and differently from the stimulating apparatus of ref.^29^, our stimulation was primarily caudo-rostral instead of ventro-dorsal, accompanied by a minimal dorso-ventral component. Thanks to the spring between the moving head and the fixed body of the miniature solenoid, de-electrifying it provoked the elastic return of the plunger and of the whiskers group to their resting position. Electrical and hardware components serving the mechanics were an Ardunio board (Genuino Uno), a variable power supplier (0-15 V, 0-3 A; LABPS1503, VALLEMAN), 2 dual motor drivers (TB6612FNG, Sparkfun), and 4 relays (5 V; G46, FINDER). Each of the motor drivers could electrify up to 2 miniature solenoids (for a maximum of 4 usable), and their standby state together with the selection of the active solenoid were controlled by Arduino. When active (standby off), the motor drivers allowed a 15 ms passage of current from the power supply directed in parallel towards the solenoid and its associated relay. The relays were used to collect the triggers accompanying the stimulation through the CED1401, thus aligned to the ongoing neural activity in Spike2. To end the stimulation, the passage of current was interrupted by deactivating the motor driver (standby on). In each experiment, the current was fixed at 7.3 V to obtain the maximal performance of the miniature solenoids. At this voltage and measured via a digital force gauge (resolution 0.001 N; FH2, SAUTER GmbH), the solenoids could exert a longitudinal pulling force of 7.4 ± 1.8 mN (mean ± std) on the whiskers group, provoking an abrupt displacement. To compensate for possible light differences in pulling force or ringing across solenoids, they were routinely and randomly assigned to different whiskers groups (i.e., contra- or ipsilateral, row A or row E stimulations) across experiments. **(b)** Example of row A-whiskers stimulation before (left) and after (right) retraction of the stimulating wire by the solenoid. White dashed lines indicate the position of the whiskers group. Red arrow indicates the direction of whiskers displacement. **(c)** Example LFP traces of a cortical somatotopic mapping (row A; see Methods) indicating responses outside the intended cortical whiskers representation (out) and over it (in). Vertical bar= 200 μV, horizontal bar= 10 ms. **(d)**. Correction for recording depth in amBC. Display readings (grey) were compared with anatomical recovery of the neuron (blue). We applied the formula: D = d*cos (30°), where D is the adjusted depth and d is the depth given by the display reading. Since in this manner we could minimally reduce the discrepancy of recording depth in these 6 randomly chosen brains with recovered neurons in amBC, we applied the same correction to remaining amBC neurons.

## SUPPLEMENTARY INFORMATION: TABLES

**Tab. 1.**
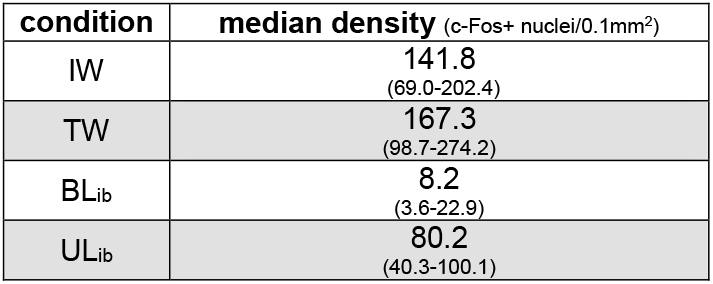
Density of c-Fos expression in the barrel cortex in the different conditions of whisker-pads intervention. Values are expressed ad median (25^th^-75^th^ quantiles). IW= intact whiskers; TW= trimmed whiskers; BL_ib_= bilateral input block; UL_ib_= unilateral input block.

**Tab.2.**
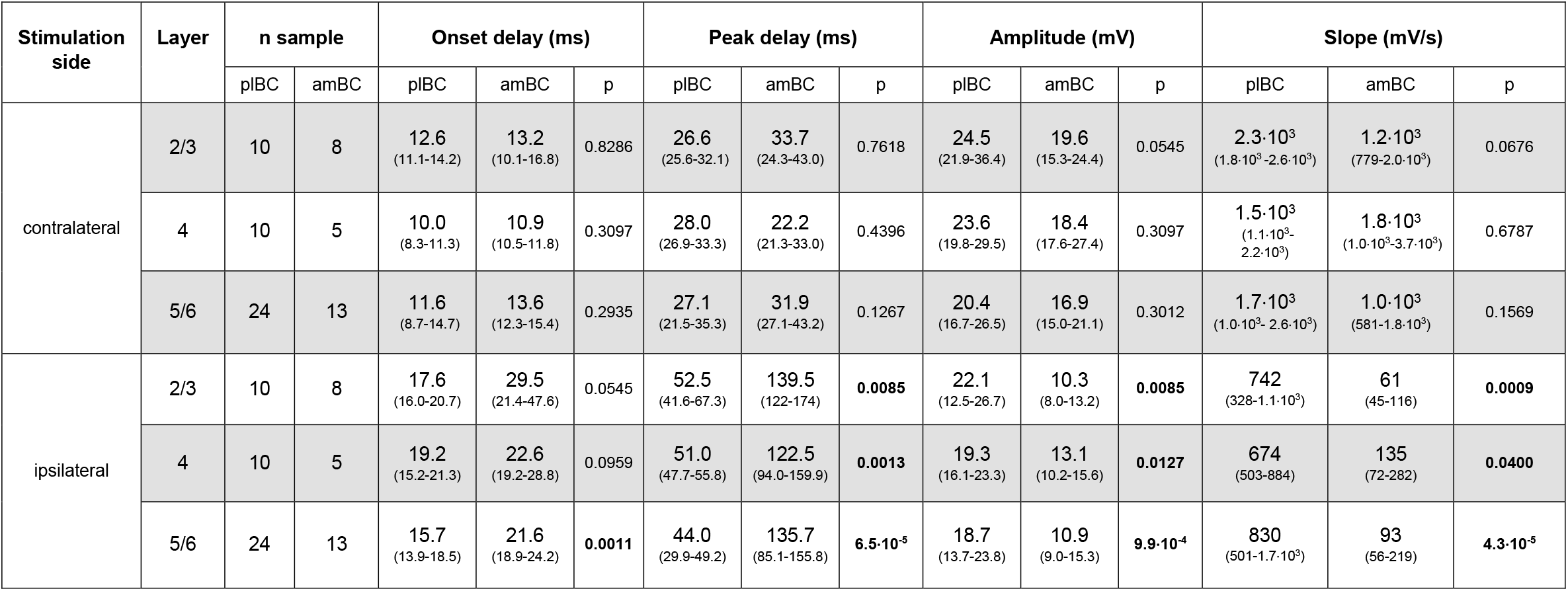
Statistical comparisons between amBC and plBC of contra- and ipsilateral whisker responses in different layers of the barrel cortex. Contra- and ipsilateral stimulations of row A- and row E-whiskers evoke in the same neurons similar contra-but different ipsilateral responses between plBC and amBC. Data are shown as median (25^th^-75^th^) quantiles. Significant p-values of the Wilcoxon rank sum test are highlighted in bold.

**Supplementary Tab.3.** Statistical comparison of membrane properties between plBC and amBC neurons. Electrophysiological membrane properties in neurons do not differ in the two BC subregions. Data are expressed as median (25^th^-75^th^) quantiles.

| Parameter | BC subregion |  | Wilcoxon rank sum test |
| --- | --- | --- | --- |
|  | pIBC | amBC | p |
| $\tau$ (ms) | 8.0<br>(2.3-10.9) | 6.9<br>(4.7-17.1) | 0.4365 |
| $R_{\text{down}}$ (MΩ) | 51.0<br>(42.1-65.6) | 49.9<br>(41.1-76.3) | 0.8506 |
| $R_{\text{up}}$ (MΩ) | 49.9<br>(43.6-62.2) | 47.8<br>(38.5-69.7) | 0.7154 |
| $C_{\text{down}}$ (pF) | 133.4<br>(54.5-225.8) | 133.2<br>(92.8-189.8) | 0.8126 |
| $C_{\text{up}}$ (pF) | 137.3<br>(55.3-223.1) | 136.5<br>(103.2-210.9) | 0.5555 |
| $V_{\text{mrest}}$ (mV) | -64.9<br>(-72.2 -56.8) | -62.5<br>(-70.8 -55.4) | 0.3915 |

**Supplementary Tab.4.**
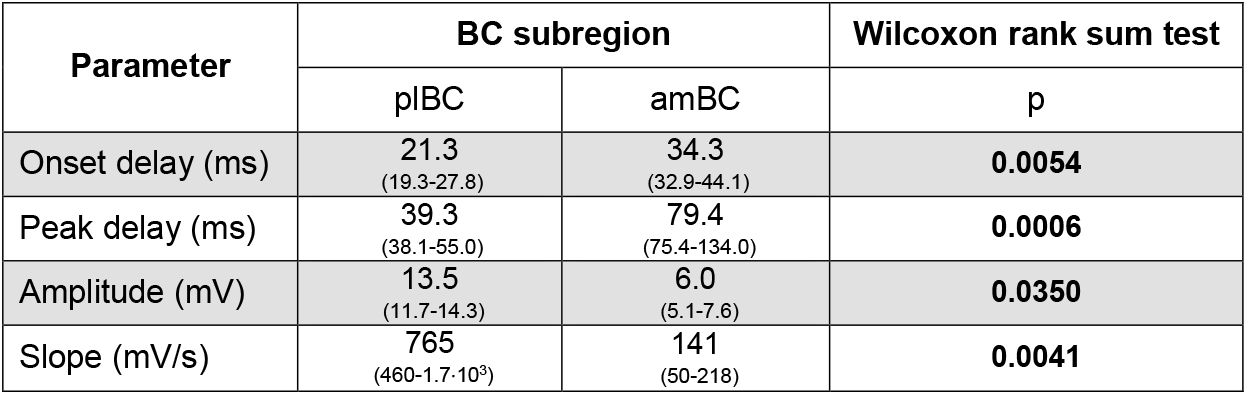
Statistical comparison of inhibitory response properties between plBC and amBC neurons. Data demonstrate the presence of a stronger synaptic recruitment in the posterolateral barrel cortex than in the anteromedial one. Data are expressed as median (25^th^-75^th^) quantiles. Significant p-values are provided in bold.

| Parameter | BC subregion |  | Wilcoxon rank sum test |
| --- | --- | --- | --- |
|  | pIBC | amBC | p |
| Onset delay (ms) | 21.3<br>(19.3-27.8) | 34.3<br>(32.9-44.1) | **0.0054** |
| Peak delay (ms) | 39.3<br>(38.1-55.0) | 79.4<br>(75.4-134.0) | **0.0006** |
| Amplitude (mV) | 13.5<br>(11.7-14.3) | 6.0<br>(5.1-7.6) | **0.0350** |
| Slope (mV/s) | 765<br>(460-1.7·10<sup>3</sup>) | 141<br>(50-218) | **0.0041** |

**Supplementary Tab. 5.**
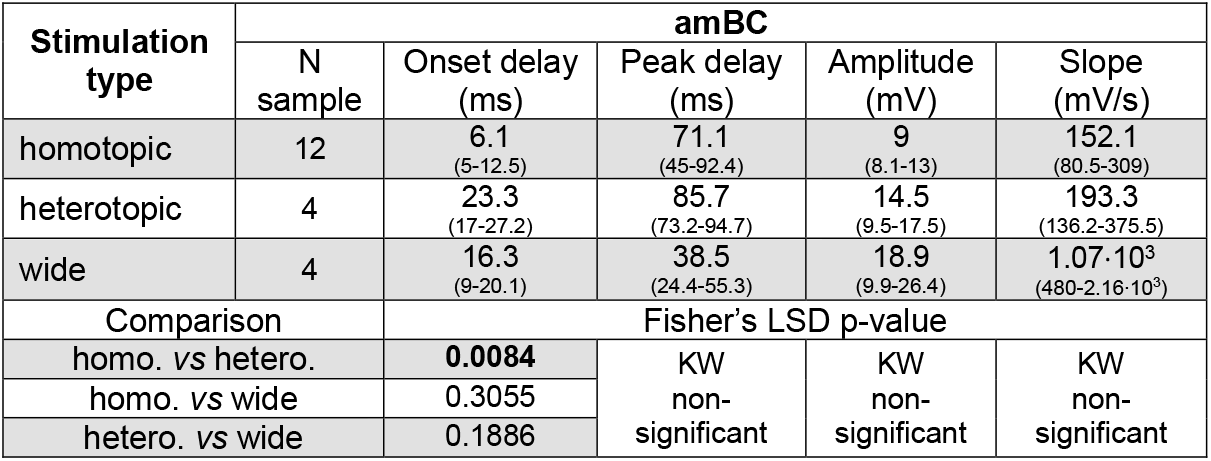
Statistical comparison of response parameters to homo-, heterotopic and wide optogenetics stimulation in amBC neurons. N samples refers to the number of neurons. KW refers to the Kruskal-Wallis test. Data are expressed as median (25^th^-75^th^) quantiles. Significant p-values are provided in bold.

| Stimulation type | amBC |  |  |  |  |
| --- | --- | --- | --- | --- | --- |
|  | N sample | Onset delay (ms) | Peak delay (ms) | Amplitude (mV) | Slope (mV/s) |
| homotopic | 12 | 6.1<br>(5-12.5) | 71.1<br>(45-92.4) | 9<br>(8.1-13) | 152.1<br>(80.5-309) |
| heterotopic | 4 | 23.3<br>(17-27.2) | 85.7<br>(73.2-94.7) | 14.5<br>(9.5-17.5) | 193.3<br>(136.2-375.5) |
| wide | 4 | 16.3<br>(9-20.1) | 38.5<br>(24.4-55.3) | 18.9<br>(9.9-26.4) | 1.07·10<sup>3</sup><br>(480-2.16·10<sup>3</sup>) |
| Comparison |  | Fisher's LSD p-value |  |  |  |
| homo. vs hetero. |  | **0.0084** | KW non-significant | KW non-significant | KW non-significant |
| homo. vs wide |  | 0.3055 |  |  |  |
| hetero. vs wide |  | 0.1886 |  |  |  |

**Supplementary Tab. 6.**
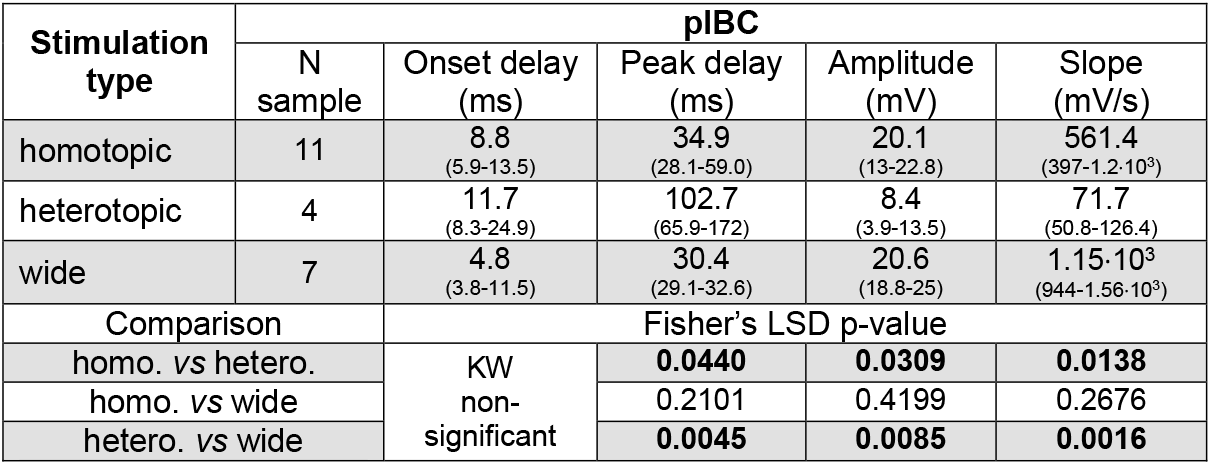
Statistical comparison of response parameters to homo-, heterotopic and wide optogenetics stimulation in pIBC neurons. N samples refers to the number of neurons. KW refers to the Kruskal-Wallis test. Data are expressed as median (25^th^-75^th^) quantiles. Significant p-values are provided in bold.

| Stimulation type | pIBC |  |  |  |  |
| --- | --- | --- | --- | --- | --- |
|  | N sample | Onset delay (ms) | Peak delay (ms) | Amplitude (mV) | Slope (mV/s) |
| homotopic | 11 | 8.8<br>(5.9-13.5) | 34.9<br>(28.1-59.0) | 20.1<br>(13-22.8) | 561.4<br>(397-1.2·10<sup>3</sup>) |
| heterotopic | 4 | 11.7<br>(8.3-24.9) | 102.7<br>(65.9-172) | 8.4<br>(3.9-13.5) | 71.7<br>(50.8-126.4) |
| wide | 7 | 4.8<br>(3.8-11.5) | 30.4<br>(29.1-32.6) | 20.6<br>(18.8-25) | 1.15·10<sup>3</sup><br>(944-1.56·10<sup>3</sup>) |
| Comparison |  | Fisher's LSD p-value |  |  |  |
| homo. vs hetero. |  | KW non-significant | 0.0440 | 0.0309 | 0.0138 |
| homo. vs wide |  |  | 0.2101 | 0.4199 | 0.2676 |
| hetero. vs wide |  |  | 0.0045 | 0.0085 | 0.0016 |

